# Competition between transcription and loop extrusion modulates promoter and enhancer dynamics

**DOI:** 10.1101/2023.04.25.538222

**Authors:** Angeliki Platania, Cathie Erb, Mariano Barbieri, Bastien Molcrette, Erwan Grandgirard, Marit AC de Kort, Karen Meaburn, Tiegh Taylor, Virlana M Shchuka, Silvia Kocanova, Guilherme Monteiro Oliveira, Jennifer A Mitchell, Evi Soutoglou, Tineke L Lenstra, Nacho Molina, Argyris Papantonis, Kerstin Bystricky, Tom Sexton

**Affiliations:** Department of Functional Genomics and Cancer, Institute of Genetics and Molecular and Cellular Biology (IGBMC), UMR7104, Centre National de la Recherche Scientifique, U1258, Institut National de la Santé et de la Recherche Médicale, University of Strasbourg, 6704 Illkirch, France; Translational Epigenetics Group, Institute of Pathology, University Medical Centre Göttingen, Göttingen, Germany; Division of Gene Regulation, the Netherlands Cancer Institute, Oncode Institute, 1066CX Amsterdam, the Netherlands; Genome Damage and Stability Centre, Sussex University, School of Life Sciences, University of Sussex, Brighton, UK; Department of Cell and Systems Biology, University of Toronto, Toronto, Ontario M55 3G5, Canada; Molecular Cellular and Developmental Biology unit (MCD), Centre de Biologie Integrative (CBI) University of Toulouse Paul Sabatier, CNRS, 31062 Toulouse, France; Institut Universitaire de France (IUF).

**Keywords:** Chromatin dynamics, enhancer, transcription, anomalous diffusion, molecular dynamics, live imaging

## Abstract

The spatiotemporal configuration of genes with distal regulatory elements, and the impact of chromatin mobility on transcription, remain unclear. Loop extrusion is an attractive model for bringing genetic elements together, but how this functionally interacts with transcription is also largely unknown. We combine live tracking of genomic loci and nascent transcripts with molecular dynamics simulations to assess the 4D arrangement of the *Sox2* gene and its enhancer, in response to a battery of perturbations. We find that alterations in chromatin mobility, not promoter-enhancer distance, is more informative about transcriptional status. Active elements display more constrained mobility, consistent with confinement within specialized nuclear sites, and alterations in enhancer mobility distinguish poised from transcribing alleles. Strikingly, we find that whereas loop extrusion and transcription factor-mediated clustering contribute to promoter-enhancer proximity, they have antagonistic effects on chromatin dynamics. This provides an experimental framework for the underappreciated role of chromatin dynamics in genome regulation.

## INTRODUCTION

Metazoan gene expression is finely controlled by regulatory inputs from promoter sequences and distal regulatory elements, such as enhancers. How these dispersed elements are spatiotemporally coordinated to determine transcriptional output has been a topic of intense study, but key questions remain unresolved^1, 2^. For example, most chromosome conformation capture-based experiments support a model where distal enhancers require close spatial proximity to target genes to confer activation^3–5^, often presumed via chromatin “looping”, and engineered promoter-enhancer contacts have been shown to stimulate expression^6^. However, recent imaging experiments at different loci have shown clear physical separation of enhancers and active genes (often in the order of ∼200-300 nm), with conflicting results as to whether their separation decreases^7, 8^, stays the same^9^ or may actually increase^10^ on transcriptional activation. The identification of nuclear clusters of lineage-specific transcription factors^11, 12^, RNA polymerase II (PolII)^13, 14^, and genes and regulatory elements themselves^15–19^ has led to the proposal that transcription is organized into specialized nuclear microenvironments or “hubs”^20^. In this model, high local concentrations of regulatory factors accumulate near genomic sequences that are loosely co-associated; the shared location of regulatory elements within this microenvironment is more important for transcriptional regulation than their direct physical juxtaposition. These hubs are potentially built up through specific protein-protein interactions and/or phase-separated condensate formation between intrinsically disordered domains found on many transcription factors^12, 21^. How genes and distal regulatory elements locate or nucleate such hubs, and how the different regulatory inputs are coordinated into a transcriptional decision within, remain open questions.

An often-overlooked aspect determining how regulatory chromatin architectures are built or maintained is the underlying mobility (and freedom of movement) of the component factors. Whereas non-histone chromatin proteins, including transcription factors, are highly mobile in the nucleoplasm^22^, the large polymeric chromatin fiber has more constrained movement, due to a crowded environment and elastic interactions with neighboring segments of the polymer^23^. This is often characterized as *sub-diffusive* movement, whereby the mean squared displacement (MSD) of the genomic element follows the relationship with time, t: *MSD ∝ Dt^α^* where *D* is the diffusion coefficient (a proxy for diffusive speed) and *α*, the anomalous exponent, is smaller than the value of 1 found in classical Brownian diffusion. Initial studies of randomly inserted tags suggested that heterochromatin was inherently less mobile than euchromatin in yeast^24^, but we recently reported significant locus-specific differences in chromatin diffusive properties at euchromatic regions in mouse embryonic stem cells (ESCs), suggesting finer-scale modulation of chromatin dynamics, potentially linked to underlying function^25^. As methods for tracking specific genomic regions *in vivo* has recently become available, researchers have asked whether the transcriptional activity of genes affects their mobility. We previously showed that an estrogen-inducible transgene could only explore a much smaller nuclear volume on acute induction, and that this constraint was dependent on transcriptional initiation^26^, consistent with a model whereby the activated gene becomes confined within a transcriptional hub. This is supported by analogous studies showing slowed movement of the pluripotency gene *Nanog* when transcriptionally active^27^ and global enhanced chromatin mobility on treatment with transcriptional inhibitors^28, 29^. However, mobility of the *Oct4* gene was found to be insensitive to transcriptional status^27^, and a different study of transition between ESCs and epiblast-like cells reported acceleration of both promoters and enhancer-proximal sequences specific to the cell state where the elements are active^30^. A clear understanding of how transcriptional events modulate chromatin dynamics (or vice versa) is thus still lacking.

At a larger scale, promoter-enhancer communication takes place within the context of topologically associated domains (TADs), regions of enhanced intra-domain chromatin contacts identified in population-average Hi-C maps^31, 32^, which have been proposed to prevent inappropriate promoter-enhancer communication across TAD borders, and/or to reduce the effective search space for enhancers to co-associate with target genes^1, 33–35^. The major mechanism believed to organize TAD architecture is loop extrusion by SMC protein complexes, particularly cohesin (reviewed in ref. 36, and references therein). Cohesin loads on chromatin, extrudes loops of the chromatin fiber (mostly bidirectionally), and is dissociated by the factor WAPL to generate metastable chromatin loops in an equilibrium. Sites where loop extrusion are stalled, particularly the juxtaposition of convergently facing sites for the factor CTCF (CCCTC-binding factor), whose N-terminus specifically interacts with the cohesin complex^37^, form the bases of more stabilized loops at TAD borders. These are observed in Hi-C maps^38^, although recent live imaging experiments have revealed these to also be relatively rare and transient interactions in individual cells^39, 40^. The so-called “architectural interactions” brought about by convergent CTCF sites have been proposed to facilitate juxtaposition of adjacent promoters and enhancers^41^, and loop extrusion has also been shown to be required for efficient activity of more distal, as opposed to proximal, enhancers^42, 43^. Further, recent simulations and experimental results show that cohesin-mediated loop extrusion generally reduces chromatin mobility^28, 40^, perhaps further “stabilizing” such genomic configurations. However, we and others have observed extensive persistence of promoter-enhancer interactions when CTCF and/or cohesin is ablated^44–46^. This latter finding is reminiscent of the apparent competition that exists between TADs and chromosome compartmentation (higher-order co-associations between domains of homotypic chromatin, such as active with active): cohesin ablation eliminates TADs, but appears to strengthen compartments^47, 48^. The exact mechanisms promoting chromosome compartmentation genome-wide are unclear, but since co-association of active domains appears to depend on transcription itself^49^ and transcription co-factors like BRD2^50^, it is reasonable to hypothesize that these could be the same biochemical principles involved in clustering transcription factors and active elements at proposed transcriptional hubs, raising the curious possibility that loop extrusion may sometimes antagonize such nuclear microenvironments.

To assess more closely the interplay of enhancer activity, transcription and cohesin-mediated loop extrusion on the spatiotemporal organization of gene regulation, we tagged the promoter and major enhancer (SCR; *Sox2* control region) of the pluripotency gene *Sox2* in mouse ESCs, in combination with detection of nascent transcription from the same allele, for their tracking at high temporal resolution. With a battery of genetic and other experimental perturbations, we confirmed the decoupling of transcription and promoter-enhancer proximity that was previously described at this locus^9, 44, 46^, and also found a hitherto unappreciated localized effect of transcription on chromatin dynamics. Promoter kinetics are disrupted on severe transcriptional inhibition, but when comparing transcriptionally bursting and “poised” *Sox2* alleles, it is the enhancer which specifically becomes less mobile on gene firing. Most strikingly, we observed that deleting either CTCF or transcription factor binding sites induced opposing effects on localized chromatin mobility. In addition, molecular dynamics simulations were able to recapitulate these experimental findings only when competition between loop extrusion and transcription-linked enhancer-promoter communication was reinforced, suggesting they may be antagonistic processes on chromatin dynamics, even though from a population-average viewpoint they are assumed to mediate the same interaction. These findings unify some of the seemingly conflicting results from previous studies and highlight the pitfalls of relying solely on population-averaged experiments on fixed cells to draw conclusions about a dynamic process such as spatiotemporal gene regulation.

## RESULTS

### Enhancer-promoter proximity is frequently maintained at the *Sox2* locus independent of transcription

To follow promoter-enhancer communication in living cells, we tagged the *Sox2* promoter and SCR with orthologous *parS* sites (ANCHOR1 and ANCHOR3) in the *musculus* allele of F1 (*Mus musculus*^129^ × *Mus castaneus*) hybrid ESCs (we term this line *Sox2-SCR_WT_*), enabling their visualization on transfection with plasmids encoding fluorescently labeled ParB proteins^26, 51^. The small size (∼1 kb) of the *parS* sequences allowed us to place the tags very close to the promoter (5.5 kb upstream of the TSS) and actually inside the SCR, in between the elements known to control *Sox2* transcription^44^, facilitating direct imaging of the promoter and enhancer themselves. As a control, we also generated an ESC line (*Inter-Down*) with *parS* integration sites shifted by ∼60 kb, allowing visualization of non-regulatory loci with equivalent genomic separation (**Fig 1a,b**). Allele-specific qRT-PCR showed that neither integration of the *parS* sites, nor their binding by transfected ParB proteins, affected *Sox2* expression in *cis* (**Fig 1c**). Cell cycle progression and expression of other pluripotency markers was similarly unaffected by the *parS*/ParB system (**Fig S1a,b**). Further, allele-specific 4C-seq showed that *Sox2*-SCR interaction strength was also unaltered by *parS* integration and ParB binding (**Fig 1d**), strongly suggesting that the imaging setup has negligible functional impact on the loci that are visualized in this study.

**Figure 1.**
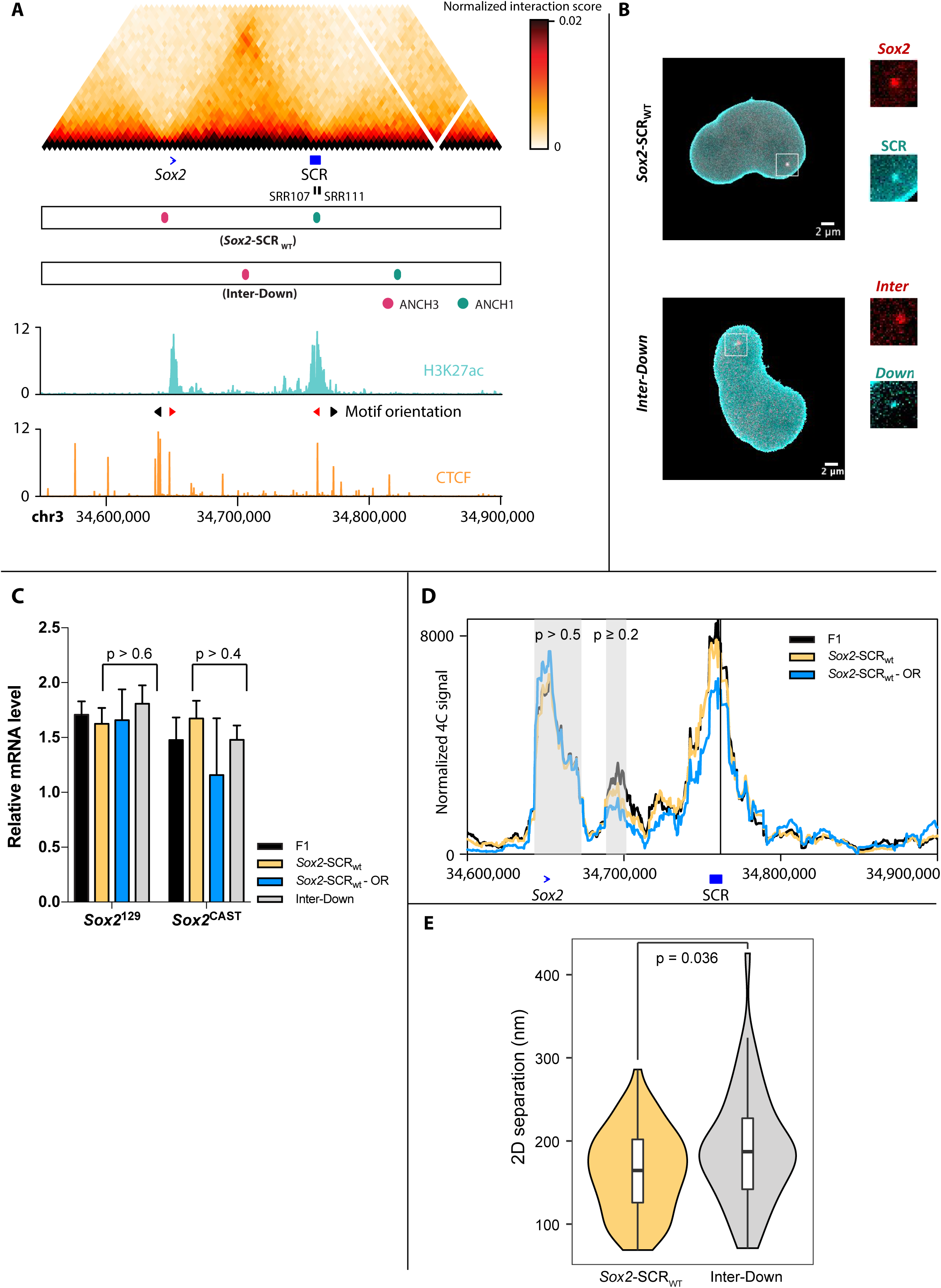
*parS*/ParB labeling of specific sites at the *Sox2* locus. **a**) Overview of the *Sox2* locus used in this study, showing (top to bottom): Hi-C map in ESCs (data from ref 60), showing the TAD delimited by the *Sox2* gene and SCR (positions shown in blue, and the locations of specific SCR regulatory elements SRR107 and SRR111 shown in black, below the map); scaled positions of ANCH1 and ANCH3 labels inserted in the *Sox2-SCR_WT_* and *Inter-Down* ESC lines; ESC ChIP-seq profiles for the active histone mark, acetylation of histone H3 lysine-27 (H3K27ac; cyan) and CTCF (orange) at the same locus (data from ref 88). Orientations of the motifs of the major CTCF sites are denoted by arrows above the CTCF ChIP-seq profile; the red arrows denote the positions of CTCF sites that are perturbed in this study. **b**) Representative images of *Sox2-SCR_WT_*(top) and *Inter-Down* (bottom) nuclei (after segmentation and removal of cytoplasmic signal), with the OR3-IRFP signal shown in red and the OR1-EGFP signal shown in cyan; scale bar 2 µm. Insets show zoomed regions around spots corresponding to bound *parS* sequences. **c**) Allele-specific qRT-PCR quantification, normalized to *SDHA*, of *Sox2* expression in F1, *Sox2-SCR_WT_* before and after transfection with ParB vectors (OR), and *Inter-Down* ESCs (*n* >= 3). Error bars indicate standard deviations from the mean. For both alleles, expression is not significantly different to that measured in F1 cells (minimum *p*-values from two-tailed t-tests are given). **d**) Musculus-specific 4C-seq profiles (mean of two replicates) using the SCR as bait are shown for F1 and *Sox2-SCR_WT_* ESCs, before and after transfection with ParB vectors (OR). The two regions consistently called as interactions are denoted in gray, and the minimum *p*-values for compared interaction scores with F1 (two-tailed t-test) are denoted. **e**) Violin plot showing the distributions of median inter-probe distances for *Sox2-SCR_WT_* and *Inter-Down* ESCs. *Sox2* and SCR are significantly closer on average than control sequences (*p* = 0.036; Wilcoxon rank sum test).

For these two ESC lines, we imaged the labeled loci at high temporal resolution (500 ms per frame) in 2D with spinning disk confocal microscopy, using a set of optimized affine transformations to correct for chromatic aberrations and control for camera misalignment. Further enhancement of spot centroid localization was performed by fitting a 2D Gaussian shape to a precision of ∼30 nm^25^. In line with previous high-throughput DNA FISH studies^52, 53^, median inter-probe distances demonstrated a large cell-to-cell heterogeneity, varying from 69-286 nm (*n* = 72) for *Sox2*-SCR distances and from 71-426 nm (*n* = 49) for the control regions (**Fig 1e**; **Table S1**). Over the short periods over which images were acquired, separation distances were relatively stable (median absolute deviation 30-107 nm; inter-decile range 74-293 nm). As may be expected from ESC Hi-C maps, the median distance between *Sox2* and the SCR (164 nm) is significantly smaller than the control regions of equivalent genomic separation (187 nm; *p* = 0.036; Wilcoxon rank sum test). The *Sox2*-SCR separation distances we measured agree very well with the 2D distances measured in the wild-type *Sox2* locus in recent multiplexed DNA FISH studies^53^ (median 160 nm; *p* = 0.78; Wilcoxon rank sum test) (**Fig S2a**). Although this implies that the *parS*/ParB system does not perturb endogenous chromosomal locus configurations within the nucleus, the ANCHOR1-tagged control region downstream of the SCR was not included in the multiplexed DNA FISH design. We therefore performed 3D DNA FISH in wild-type F1 and ANCHOR-labeled *Sox2-SCR_WT_*ESCs, using fosmid probes covering the locations of each of the *parS* integration sites, further confirming that average *Sox2*-SCR separation distances are smaller than the control regions (*p* < 2.2×10^-16^; Wilcoxon rank sum test), and that ANCHOR labeling has negligible effects on underlying chromatin organization (*p* = 0.14; Wilcoxon rank sum test) (**Fig S2b,c**). We note that a previous study using Tet-and Cu-repeat operators to label regions a little further upstream of the *Sox2* promoter and downstream of the SCR measured significantly larger distances (median 221 nm; *p* = 5.4×10^-8^; Wilcoxon rank sum test) than obtained with either ANCHOR or DNA FISH in untagged loci^9^ (**Fig S2a**). This suggests that, at least for this locus, the repetitive operator system may artificially inflate physical separation of tagged loci compared to untagged counterparts.

We previously identified by 4C-seq a near-complete loss of *Sox2*-SCR interaction in the early stages of ESC differentiation to neuronal precursors when *Sox2* expression is shut down^54^. We recapitulated this finding when differentiating the labeled *Sox2-SCR_WT_* cells, observing a significant increase in average *Sox2*-SCR distance (median 200 nm; *p* = 0.02; Wilcoxon rank sum test) by the third day, in agreement with the timing of loss of interaction observed by allele-specific 4C (**Fig 2a,b**). This cannot be solely explained by general chromatin compaction changes accompanying loss of pluripotency, because the average distances between the control regions were not significantly increased (**Fig 2c**; median 208 nm; *p* = 0.15; Wilcoxon rank sum test). It has been proposed that promoter-enhancer “interactions” are somehow causal in transcriptional activation^3, 6^, and the onset of increased promoter-enhancer separation corresponds to near-complete transcriptional shutdown of *Sox2* (**Fig S1c**). However, recent studies, including those at the murine *Sox2* locus, have brought this into question^9,10,44^. To directly test the effect of transcription on promoter-enhancer proximity, we also measured *Sox2*-SCR distances upon either global inhibition of transcription by treatment of the *Sox2-SCR_WT_* cells with the drug triptolide (which globally shuts down transcription by inhibiting TFIIH and inducing PolII degradation)^55^, or specific inhibition of *Sox2* expression in *cis* by the allele-specific deletion of two critical elements of the SCR (SRR107 and SRR111)^44^ to create the line *Sox2-SCR_ΔSRR107+111_*. In both cases, despite efficient loss of total or *musculus*-specific *Sox2* expression, respectively, observed with qRT-PCR and single-molecule RNA FISH (**Fig S1d-f**), average *Sox2*-SCR (and control region) distances remained small and were non-significantly altered (median distances 142-172 nm; *p* > 0.05; Wilcoxon rank sum test; **Fig 2d**; **Fig S2d**). Overall, we thus find that the *Sox2* promoter and SCR are frequently in relatively close proximity, over the sort of range traditionally called as “interactions” or “looping events” (<200 nm), and significantly closer than control regions of equivalent genomic separation. However, whereas early stages of neuronal differentiation can increase average physical separation between promoter and enhancer, their distances remain relatively close. Moreover, the promoter and enhancer remain proximal on orthogonal perturbations disrupting *Sox2* transcription to the same extent, such as deletion of key SCR elements or pharmacological inhibition of transcriptional initiation. The idea that chromatin “interaction” and transcriptional activation can be decoupled is thus further reinforced.

**Figure 2.**
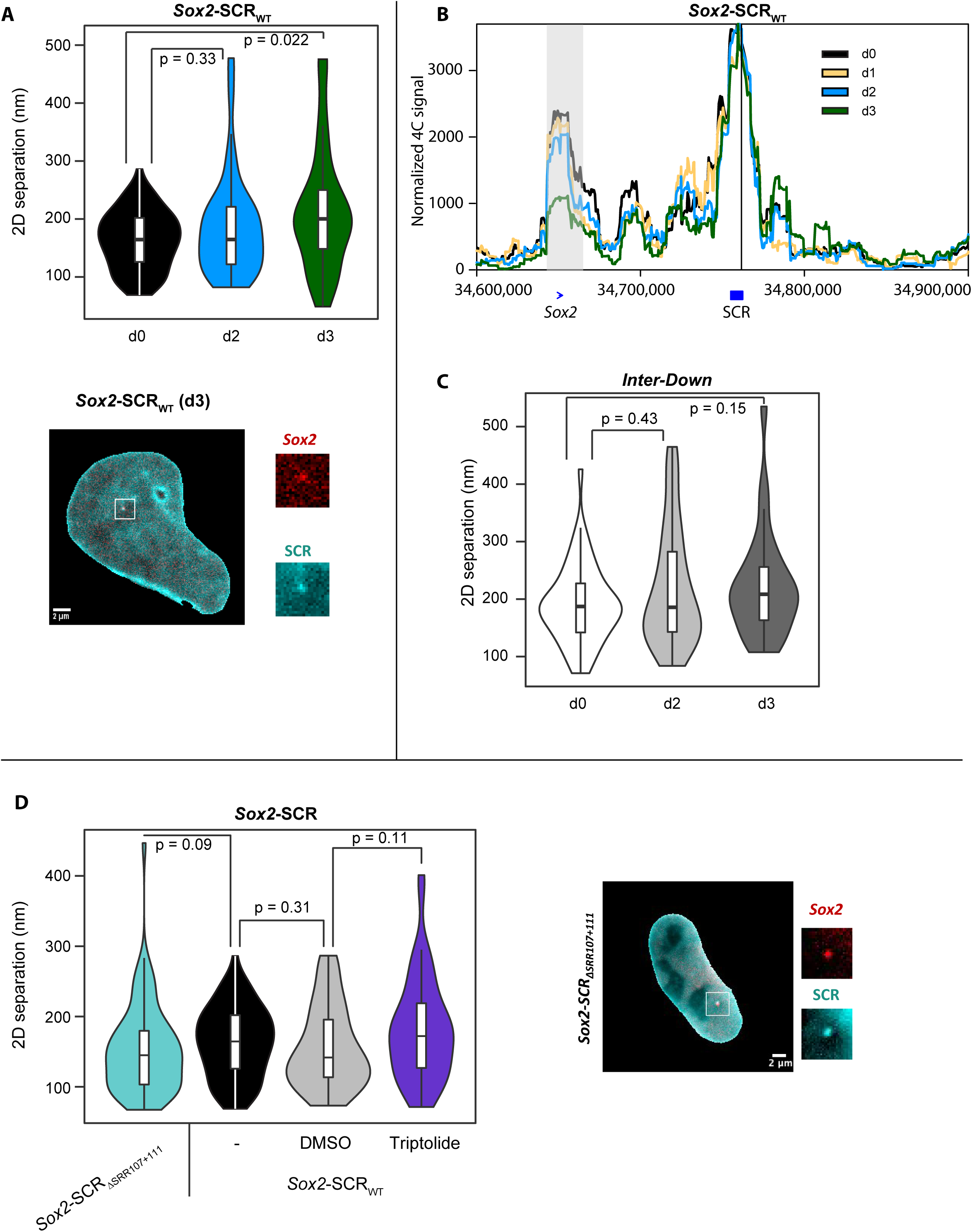
*Sox2*-SCR distance is frequently uncoupled from transcriptional status. **a**) Violin plot showing the distributions of median *Sox2*-SCR distances in *Sox2-SCR_WT_*ESCs after 0, 2 and 3 days of differentiation. Comparisons are made by Wilcoxon rank sum tests, with *p*-values given. Below is shown a representative image of a *Sox2-SCR_WT_* nucleus after 3 days of differentiation, with the OR3-IRFP signal shown in red and the OR1-EGFP signal shown in cyan; scale bar 2 µm. Insets show zoomed regions around spots corresponding to bound *parS* sequences. **b**) Musculus-specific 4C-seq profile, using the SCR as bait, for *Sox2-SCR_WT_* ESCs at 0, 1, 2 and 3 days after differentiation. A dramatic loss of *Sox2*-SCR interaction (denoted by gray region) is observed on the third day. **c**) Violin plot showing the distributions of median inter-probe distances for *Inter-Down* ESCs after 0, 2 and 3 days of differentiation. Comparisons are made by Wilcoxon rank sum tests, with *p*-values given. **d**) Violin plot showing the distributions of median inter-probe distances for *Sox2-SCR_ΔSRR107+111_* ESCs, and *Sox2-SCR_WT_* cells after no treatment (-), or treatment with DMSO or triptolide. Comparisons are made by Wilcoxon rank sum tests, with *p*-values given. To the right is shown a representative image of a *Sox2-SCR_ΔSRR107+111_* nucleus, with the OR3-IRFP signal shown in red and the OR1-EGFP signal shown in cyan; scale bar 2 µm. Insets show zoomed regions around spots corresponding to bound *parS* sequences.

### Promoters and enhancers have more constrained chromatin motion than control sequences

We previously reported that seemingly neutral elements within the HoxA locus exhibit different and locus-specific diffusive properties of labeled chromatin^25^. We therefore questioned whether chromatin mobilities are further different in promoters and enhancers, and whether these properties are linked to transcriptional output. Indeed, on examination of the tracks of labeled regions around the *Sox2* locus, as well as their ensemble MSD curves, it was apparent that the promoter and enhancer were more limited to explore the nuclear volume than the control regions (**Fig 3a,b**). To assess the local chromatin mobilities more quantitatively, we applied our previously developed GPTool to fit the track trajectories to a fractional Brownian motion (FBM) model and extract their apparent diffusion coefficients, *D_α_app_*, and anomalous exponents, *α_app_*^25^. As previously, we confirmed that the trajectories approximated to self-similar Gaussian-distributed displacements and an FBM velocity autocorrelation function over the time scale in which measurements are taken (**Figs S3** and **S4**), affirming the suitability of the model used. The experimental MSD profiles were non-linear when plotted on double-logarithmic scales, suggesting that their properties cannot be explained by one regime, likely due in the main part to the confounding effects of nuclear and cellular movement and rotation. When analyzing cross-correlations of co-measured trajectories to estimate global movements of the cell/nuclear substrate^25^, the experimental MSD curves fit very well as the sum of the component locus and substrate FBM regimes (see Methods; **Fig S5a**). We stress that GPTool, apart from assuming proximity to a global FBM regime, is actually agnostic to any specific polymer model, and obtains estimates of diffusive parameters for individual trajectories which can vary quite a lot from the average (**Fig 3c**). As such, the obtained values of the apparent anomalous exponent, *α_app_*, give a measure of constraint to the particle movement, which can be fairly compared across experiments, but do not necessarily reflect the α value derived from conventional polymer physics models for bulk diffusive properties (see also Discussion). In line with our initial observations, the average substrate-corrected apparent anomalous exponent was significantly smaller for the active elements, the *Sox2* promoter (median *α_app_* 0.130) and the SCR (median *α_app_*0.203), than for the control regions (median *α_app_* 0.276 and 0.224 for intervening and downstream regions, respectively), indicating a greater constraint to their local movements (**Fig 3c**; full data given in **Table S1**). We note that these measured values correspond well with the *α* ∼ 0.2 determined from fits to radial MSD curves for lac/TetO-tagged CTCF-mediated interacting sites with a similar genomic separation of ∼150 kb^40^. Analogous radial MSD analysis for the *Sox2-SCR_WT_* and *Inter-Down* trajectories revealed similarly constrained motion, with the promoter and enhancer significantly more constrained than the neutral sequences (*p* = 5.7×10^-6^; ANCOVA test; **Fig S5b**). However, the nature of this analysis, by comparing the motion of one locus relative to the “fixed” second locus, does not allow any mobility differences between the two loci to be discerned. Using substrate-corrected GP-FBM, we further found that the promoter exhibited significantly more constrained motion than the enhancer (*q* = 2×10^-5^; Wilcoxon rank sum test with Benjamini-Hochberg correction), and all four labeled regions around the *Sox2* locus exhibited more constrained motion than was observed around the generally inactive HoxA locus in ESCs^25^ (**Fig S5c**).

**Figure 3.**
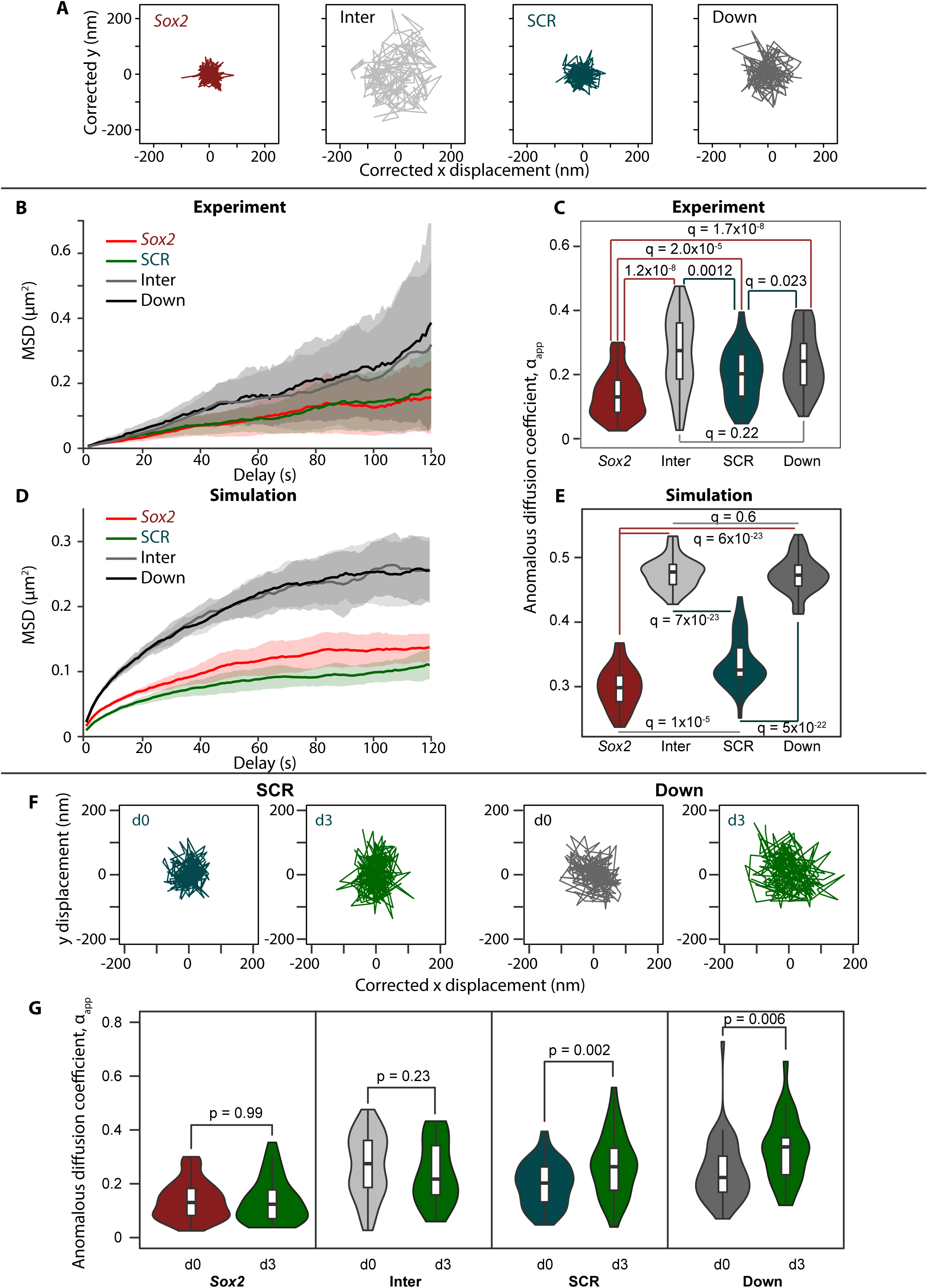
The *Sox2* promoter and SCR are more constrained than control sequences. **a**) Sample tracks of the 2D trajectories for labeled *Sox2* (red), SCR (green), Inter (light gray) and Down (dark gray) sequences, plotted on the same scale. Displacements have been corrected for substrate displacement, and the plots have been centered on the median positions. **b**) Uncorrected ensemble MSD curves for the same four regions as in **a**, with lines showing median values and shading indicating the median absolute deviation. Inter and Down elements clearly move greater distances within the same timeframe as the *Sox2* promoter and SCR. **c**) Violin plot showing the distributions of apparent anomalous exponents measured from the individual movies for each of the four regions as in **a**. Comparisons are made by Wilcoxon rank sum tests with Benjamini-Hochberg multiple testing correction, with *q*-values given. Apparent anomalous exponents are significantly lower for *Sox2* and SCR than for the control regions, demonstrating their overall greater constraint. **d**) Ensemble MSD curves for the same four regions in **a**, derived from molecular dynamics simulations of the *Sox2* locus, with lines showing median values and shading indicating the median absolute deviation. *Sox2* and SCR show apparent reduced mobility compared to control regions. **e**) Violin plot showing the distributions of apparent anomalous exponents derived from the same simulated trajectories as **d**. Comparisons are made by Wilcoxon rank sum tests with Benjamini-Hochberg multiple testing correction, with *q*-values given. **f**) Sample tracks of the 2D trajectories for labeled SCR and Down regions, at either 0 or 3 days of differentiation, plotted on the same scale. Displacements have been corrected for substrate displacement, and the plots have been centered on the median positions, showing an apparent increased freedom of movement for both regions on differentiation. **g**) Violin plots showing the distributions of apparent anomalous exponents measured from the individual movies for each of the four regions as in **a**, comparing 0 and 3 days of differentiation. Comparisons are made by Wilcoxon rank sum tests, with *p*-values given; the apparent constraint of *Sox2* and Inter is unchanged on differentiation.

These results are in line with our previous observations in a transgenic system that genes explore a more restricted nuclear volume when they are induced^26^, potentially due to their engagement in specialized nuclear microenvironments for transcription. To explore these properties further, we modeled the 800 kb of ESC chromatin around the *Sox2* locus as a self-avoiding polymer chain of 1 kb beads, classed as either “neutral”, binding PolII (*Sox2* promoter and enhancers, determined by peaks of the active histone mark, acetylation of lysine-27 of histone H3 (H3K27ac)), transcribed genic regions (*Sox2* gene body), or binding CTCF (determined by ChIP-seq peaks and with motif orientation information included). As we previously did for a theoretical locus to explain chromatin conformation changes on acute loss of PolII^56^, we performed molecular dynamics simulations using Langevin equations to model thermal motion of the chromatin and its binding factors, with the following additions to account for biological processes: PolII molecules have affinity for promoters and enhancers, and traverse the gene body to simulate transcription; PolII molecules also have affinity to each other to simulate condensate formation; cohesin complexes can bind anywhere on the polymer and extrude loops, but has preference to load at promoters and enhancers, and is unable to process past a CTCF-bound site when the motif is oriented towards the extruding loop (see Methods). The ensemble of chromatin conformations resulting from these simulations generated a contact map closely resembling that of the experimental Hi-C results (**Fig S5d**). Collecting the trajectories of the beads corresponding to the *parS*-tagged loci, we derived simulation MSD curves indicating a clear reduced mobility of *Sox2* and SCR compared to control sequences (**Fig 3d**). In accordance, application of GPTool to the simulated trajectories gave significantly smaller *α_app_* measurements at the promoter and enhancer than control regions (*q* < 5×10^-22^; Wilcoxon rank sum test with Benjamini-Hochberg correction) (**Fig 3e**). This relatively simplified depiction of the locus thus supports the experimental results, suggesting that promoter-enhancer “affinity” brought about by transcriptional regulation and/or loop extrusion processes is sufficient to explain their reduced chromatin dynamics.

### Transcriptional perturbation specifically reduces *Sox2* promoter mobility

We next measured the chromatin dynamics of the labeled loci on different means of transcriptional perturbation, reasoning that mobility constraints at the promoter and/or enhancer may be specifically relaxed. However, the distribution of apparent anomalous exponents of the *Sox2* promoter did not change when the gene was silenced by either onset of differentiation or deletion of SRR107/SRR111 (**Fig 3f,g**, **6b**), and *α_app_* of both the enhancer and promoter was only weakly and non-significantly increased on treatment with triptolide (*p* > 0.05; Wilcoxon rank sum test; **Fig S5e**). On the other hand, the SCR became significantly less constrained at the onset of ESC differentiation (median *α_app_* 0.263; *p* = 0.002; Wilcoxon rank sum test), but so did the downstream control region (median *α_app_* 0.337; *p* = 0.006), raising questions as to whether this change is directly linked to transcriptional changes (**Fig 3f,g**). Previous imaging studies, including our own, suggest that the dynamics of other gene loci are altered on their induction^26, 27, 30^, prompting us to look closer at the *Sox2* locus trajectories. Indeed, inspection of the substrate movement-corrected ensemble MSD curves revealed that promoter mobility is generally reduced when transcription is blocked by either triptolide treatment or deletion of SRR107/SRR111 (**Fig 4a,b**). The other parameter commonly used to quantify diffusive motion is the diffusion coefficient, *D_α_*, but since this is expressed in units of µm^2^/s^α^ (i.e. scaling according to *α*), its measurement can only be fairly compared between sets of trajectories containing exactly the same anomalous exponent. To quantify *Sox2* promoter dynamics while accounting for the clear variability in measured *α_app_*between individual trajectories within a population (**Fig 3c**), we used the *D_α_app_*and *α_app_* values measured for each trajectory and computed *r*, the expected radius that the locus can explore in a given time interval, after accounting for substrate movement (see Methods). For all timescales appropriate for our imaging conditions (0.5-60 s), the *Sox2* promoter explored significantly less area when transcription was perturbed than during control conditions (*q* < 0.05; Wilcoxon rank sum test with Benjamini-Hochberg multiple testing correction; **Fig 4c**). It was previously proposed that “ongoing transcription may provide a source of nonthermal molecular agitation that can “stir” the chromatin within the local chromosomal domain”^30^. Our results partially support this hypothesis, although triptolide treatment had negligible effects on SCR dynamics (**Fig 4d**), despite this enhancer producing transcripts in ESCs^57^. Further, general chromatin mobility, as measured in tagged histones, was found to actually be increased on pharmacological inhibition of transcription^28^. Taken together, our results suggest a clear but complex contribution of gene activity to chromatin dynamics. On the one hand, active elements are significantly more constrained than “neutral” genomic sequences, but transcriptional inhibition does not appear to “rescue” chromatin mobility at the former sequences. Conversely, we find evidence that a highly constrained promoter becomes even less mobile at extreme perturbation of target gene expression.

**Figure 4.**
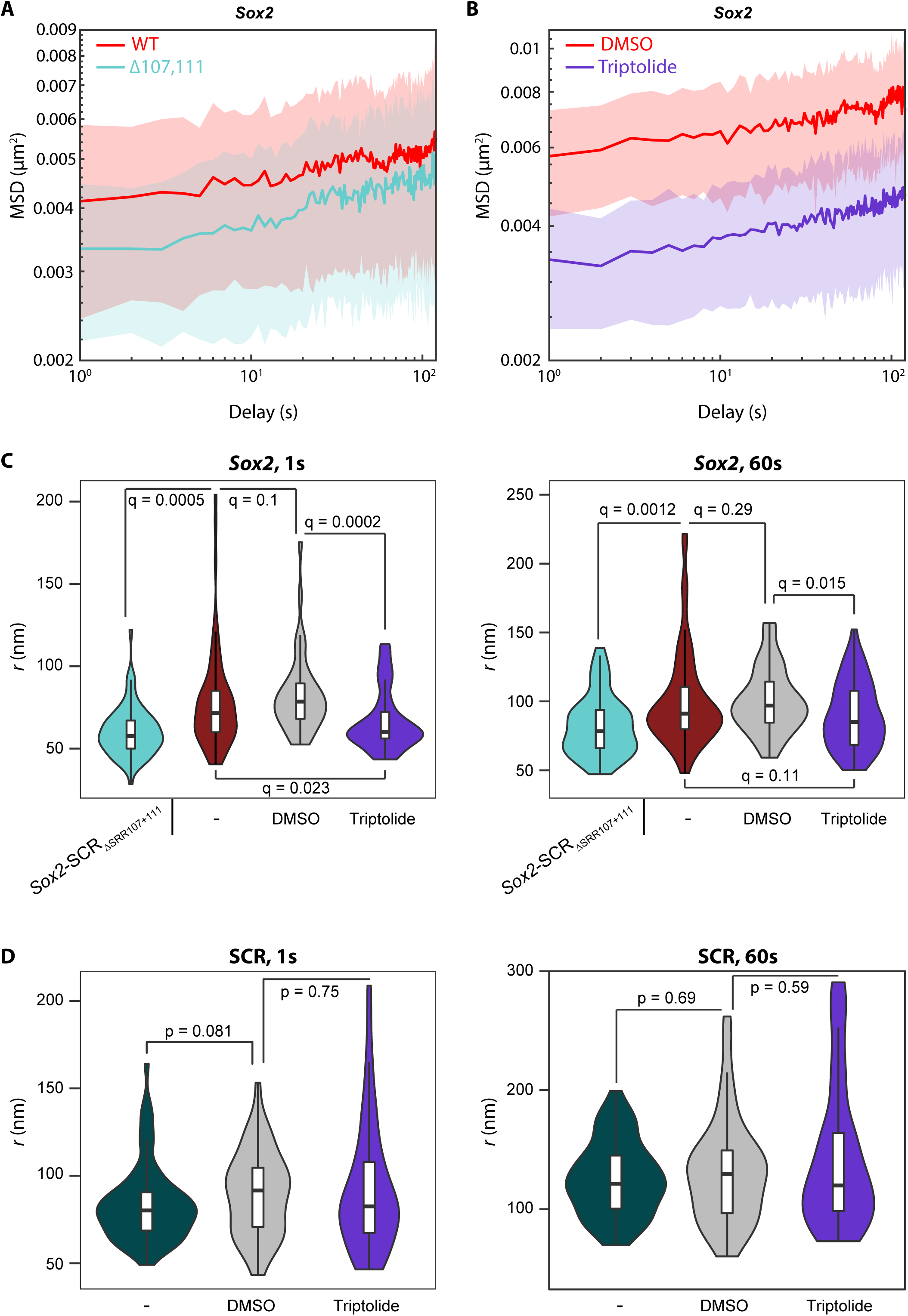
Severe transcriptional inhibition specifically reduces *Sox2* promoter mobility. **a,b**) MSD ensemble curves, corrected for substrate movement and plotted on a double-log scale, comparing *Sox2* promoter dynamics between **a**) *Sox2-SCRWT* and *Sox2-SCRΔSRR107+111* ESCs, and **b**) *Sox2-SCRWT* cells treated with either DMSO or triptolide. Lines indicate median values and shading the median absolute deviations. MSD reduction as shown by downwards vertical shift of profile on transcriptional inhibition demonstrates a reduced mobility of the labeled promoter. **c,d**) Violin plots showing distributions of apparent explored radii after 1 s (left) or 60 s (right), computed from individual movies for **c**) *Sox2* and **d**) SCR, comparing *Sox2-SCR_ΔSRR107+111_* ESCs, and *Sox2-SCR_WT_* cells after no treatment (-), or treatment with DMSO or triptolide. Comparisons are made by Wilcoxon rank sum tests with Benjamini-Hochberg multiple testing correction, with *q*-values given. *Sox2*, but not SCR, explored radii are consistently reduced on transcriptional inhibition.

### The SCR has reduced chromatin dynamics on transcriptional firing

The experiments performed so far have relied on perturbations, such as cell differentiation, *cis*-sequence deletions or treatment with drugs, to study effects of transcription, which are likely to have other indirect and confounding effects. To formally compare chromatin dynamics at transcribing versus non-transcribing *Sox2* alleles, we generated triple-labeled ESCs (*Sox2-SCR_MS2_*), combining ANCHOR tags at the SCR and *Sox2* promoter with musculus-specific incorporation of MS2 repeats into the 3’UTR of the intronless *Sox2* gene^58^, and subsequent PiggyBac-mediated transposition of constructs for both fluorescently-labeled ParB proteins and MCP. This setup allows for simultaneous tracking of promoter, enhancer and nascent RNA (**Fig 5a,b**). Previous labeling experiments at the *Sox2* locus reported a significant reduction in transcription and SOX2 protein levels on incorporation of the MS2/MCP system^9^, presumably due to interference at the 3’UTR. We screened numerous clones to find one with the minimal perturbation of *Sox2* expression, estimating from allele-specific qRT-PCR, single-molecule RNA FISH and western blotting to have a mild reduction (∼25%) of total protein levels or estimated transcription rates in our assayed cell line, despite an apparent destabilization of the MS2-tagged mRNA when bound by MCP (**Fig S6a-c**). Imaging transcription bursts over two-hour windows, we measure an average bursting time of ∼200 s and an average time between bursts of ∼630 s. As a population, the observed cells spent ∼36% of their time with a detected burst of transcription, nearly ten times more frequently than what was observed in the previous study using a cell line with more extensively perturbed *Sox2* expression^9^. To measure chromatin dynamics at maximum temporal resolution, we then used this line for triple-label tracking over five-minute windows. Over this timeframe, the majority of movies were either always transcribing or always silent, but transition points were occasionally observed (**Fig 5b**). When comparing transcribing and non-transcribing alleles, there were negligible differences in median promoter-enhancer separation (182 nm for transcribing, 175 nm for non-transcribing; *p* = 0.6; Wilcoxon rank sum test; **Fig 5c**) or apparent anomalous exponent of the *Sox2* promoter (median 0.134 for transcribing, 0.111 for non-transcribing; *p* = 0.37; Wilcoxon rank sum test; **Fig 5d**; full data given in **Table S2**). Strikingly, however, we found that the enhancer is specifically more constrained on average when transcriptionally active (median *α_app_* 0.148 for transcribing, 0.191 for non-transcribing; *p* = 0.0078; Wilcoxon rank sum test). Moreover, whereas the *Sox2* promoter has a significantly smaller median apparent anomalous exponent than that of the SCR in non-transcribing loci (*p* = 3.1×10^-5^; Wilcoxon rank sum test), in line with what was measured in *Sox2-SCR_WT_* cells (**Fig 3c**), there is no significant difference in *α_app_* between the two elements in transcriptionally bursting alleles (*p* = 0.49; Wilcoxon rank sum test). Finally, the corrected MSD curves for transcriptionally active loci specifically display a plateau at larger time lapses (**Fig 5e** and **Fig S6d**). This is more pronounced for SCR dynamics, but the *Sox2* promoter also shows similar behavior, suggesting that both elements are effectively confined within a defined volume during transcription. It is interesting to speculate that this locus restriction is a consequence of transcription taking place in very specified nuclear foci^15^ and/or condensates^21^, although chromatin dynamics will need to be traced over longer time periods, spanning several transcription bursts, to determine whether any volume restriction is exclusive to active states. Overall, this direct observation of chromatin dynamics relative to ongoing transcription shows an expectedly high cell-to-cell variability^26^, with the recurrent trend that in wild-type ESCs, the highly expressed *Sox2* gene appears to have high constraints to local mobility that are not altered by transcriptional state. On the other hand, the SCR, while maintaining relative proximity (but not necessarily complete juxtaposition) to the target gene, has greater freedom of movement when the gene is transcriptionally poised yet inactive, but becomes constrained to levels comparable with the gene itself on transcriptional firing.

**Figure 5.**
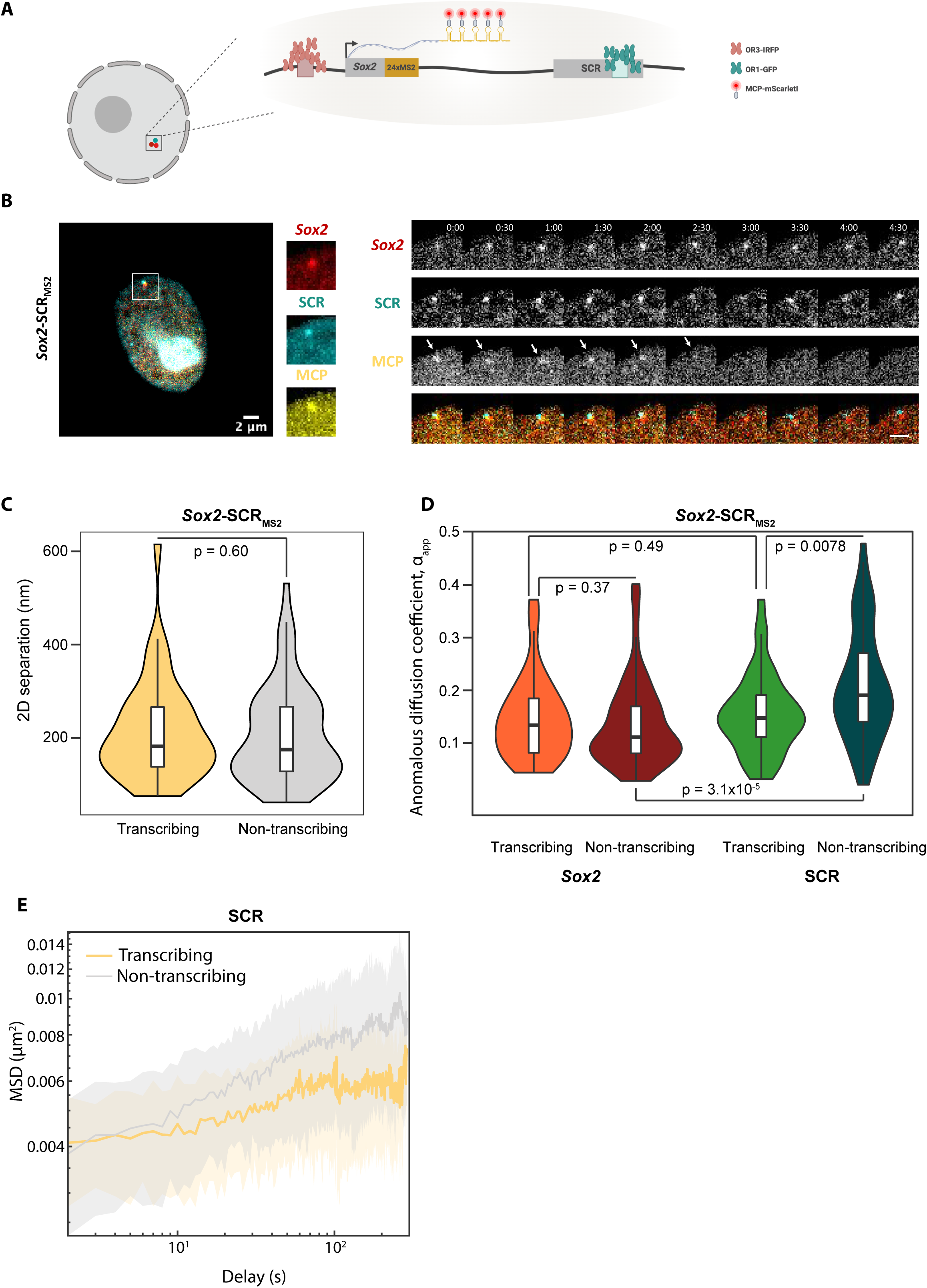
The SCR specifically becomes more constrained on transcriptional firing in wild-type loci. **a**) Schematic of triple-label system of *Sox2-SCR_MS2_* ESCs. The *Sox2* promoter DNA is tagged with ANCH3 for visualization with the ParB protein OR3-EGFP, and the SCR is tagged with ANCH1 for visualization with the ParB protein OR1-IRFP. The 3’UTR of *Sox2* contains 24 copies of the MS2 repeat for visualization of nascent RNA with MCP-ScarletI. **b**) Left: representative image of transcriptionally active *Sox2-SCR_MS2_* nucleus, with the OR3-IRFP signal shown in red, the OR1-EGFP signal shown in cyan, and the mScarletI signal shown in yellow; scale bar 2 µm. Insets show zoomed regions around spots corresponding to bound *parS* sequences and transcription site. Right: Images of the same *Sox2-SCR_MS2_*nucleus at 30 s intervals, showing each of the three channels individually in monochrome, and the overlaid image with the same color scheme as on the left. Arrows show the position of a transcription site (MCP+), which is inactivated by ∼ 3 min. Scale bar 2 µm. **c**) Violin plot showing the distributions of median inter-probe distances for transcriptionally active (MCP+) and inactive (MCP-) *Sox2-SCR_MS2_* ESCs, with no significant difference (*p* = 0.6; Wilcoxon rank sum test). **d**) Violin plot showing the distributions of apparent anomalous exponents measured from the individual movies for *Sox2* and SCR in *Sox2-SCR_MS2_* cells, comparing transcriptionally active and inactive alleles. Comparisons are made by Wilcoxon rank sum tests, with *p*-values given. **e**) MSD ensemble curves, corrected for substrate movement and plotted on a double-log scale, comparing SCR dynamics between transcriptionally active and inactive alleles. Lines indicate median values and shading the median absolute deviation. Transcribing alleles show a more distinctive plateau on the MSD curve, suggesting that there is a stricter limit on the area that is explored.

### Opposing effects of loop extrusion and promoter-enhancer communication on local chromatin mobility

Aside from transcription, chromatin topology itself may have an influence on local dynamics. For example, multiple distal interactions such as those defining TAD borders may be expected to “tether” different chromatin regions and thus impose greater constraints on their movement^59^. Differentiation to neural precursors causes loss of the ESC-specific TAD border at the SCR, whereas the strong TAD border just upstream of the *Sox2* promoter is maintained^60^ (**Fig S7a**). This loss of “anchoring” to distal regions could thus be responsible for the previously observed reduced mobility constraints at the SCR (and into the downstream TAD; **Fig 3f,g**), potentially independent of transcriptional changes. We previously showed that in ESCs, the SCR TAD border is resilient to large genetic deletions, including SRR107, SRR111 and the predominant CTCF site at the enhancer, even though the latter site was able to mediate chromatin interactions in other contexts, presumably by stalling loop extrusion^44^ (CTCF_SCR_ site in **Fig 6a**). To assess the potential effect of loop extrusion processes on local chromatin dynamics, we generated derivative lines of *Sox2-SCR_WT_* cells, harboring homozygous deletions of 16-24 nt at the core CTCF binding motifs located either just upstream of the *Sox2* promoter (*Sox2-SCR_ΔCTCF_Sox2_)* or within the SCR (*Sox2-SCR_ΔCTCF_SCR_*) (**Figs 6a** and **S7b**). As expected, ChIP-qPCR confirmed that these deletions eliminated local CTCF binding (**Fig S7c**). Both lines maintained high expression of pluripotency marker genes (**Fig S1b**), and *Sox2* expression was only mildly (and non-significantly) reduced in the *Sox2-SCR*_ΔCTCF_SCR_ line, in agreement with previous findings for homozygous deletions of this element^61^, although expression was reduced to ∼60% of wild-type levels in the *Sox2-SCR*_ΔCTCF_Sox2_ line (**Fig S7d**). In agreement with previous studies of the respective roles of these CTCF sites on *Sox2*-SCR interaction^44, 46, 61^, allele-specific 4C-seq and ANCHOR inter-probe distance measurements showed negligible topological changes when the SCR CTCF site is deleted and a mild reduced interaction on homozygous deletion of the *Sox2* CTCF site (**Fig S7e,f**). Despite these somewhat limited phenotypic effects, both lines demonstrated a significantly reduced *local* constraint on chromatin mobility when CTCF binding was abrogated: the median apparent anomalous exponent was increased at the *Sox2* promoter in *Sox2-SCR_ΔCTCF_Sox2_* cells (0.156; *p* = 0.018; Wilcoxon rank sum test) and at the SCR in *Sox2-SCR_ΔCTCF_SCR_*cells (0.235; *p* = 0.036; Wilcoxon rank sum test), with negligible, insignificant changes to *α_app_* at the regulatory element which maintains CTCF binding (**Fig 6b-e**). These findings mirror the observed increase in radial MSD on deletion of CTCF binding sites within a transgenic CTCF-mediated looping construct^40^, although the nature of this experimental setup could not distinguish whether dynamics are specifically altered locally at the CTCF anchor site, or whether they are also propagated to looping anchor partners. Our results support a more localized effect on chromatin dynamics. Previously published simulations suggest that, at short time scales consistent with our measurements, active loop extrusion increases mobility of the underlying chromatin fiber^62^, although cohesin loading itself generally imposes constraints on the underlying chromatin at larger scales^28, 40^. We propose that removal of CTCF from one site, by removing the barrier to loop extrusion through it, permits a localized relaxation of movement constraints, explaining the focal increase in *α_app_*. The observed relaxation does not appear to efficiently propagate to their “looping partners”, presumably because actual CTCF-CTCF juxtapositions are rather rare events^39^. An alternative hypothesis is that the general binding of non-nucleosomal factors to DNA changes its biophysical properties to result in “stiffer”, more constrained chromatin motion. However, this is not supported in our experiments since the median apparent anomalous exponent at the SCR is actually significantly *reduced* (i.e. even greater mobility constraints are imposed) on deletion of multiple transcription factor binding sites at SRR107 and SRR111 (0.161; *p* = 0.0048; Wilcoxon rank sum test; **Fig 6d,e**). Dispersed transcription factor binding sites at the *Sox2* and other loci have been shown to cluster via protein-protein interactions in ESCs^12^, and such homotypic interactions are believed to be a basis for chromosome compartment formation^63^. Various studies suggest that loop extrusion and chromosome compartmentation are antagonistic processes organizing population-averaged chromatin architectures^47, 48, 56, 62, 64^. It is therefore possible that processes responsible for compartmentation are also generally antagonistic to cohesin-mediated loop extrusion.

**Figure 6.**
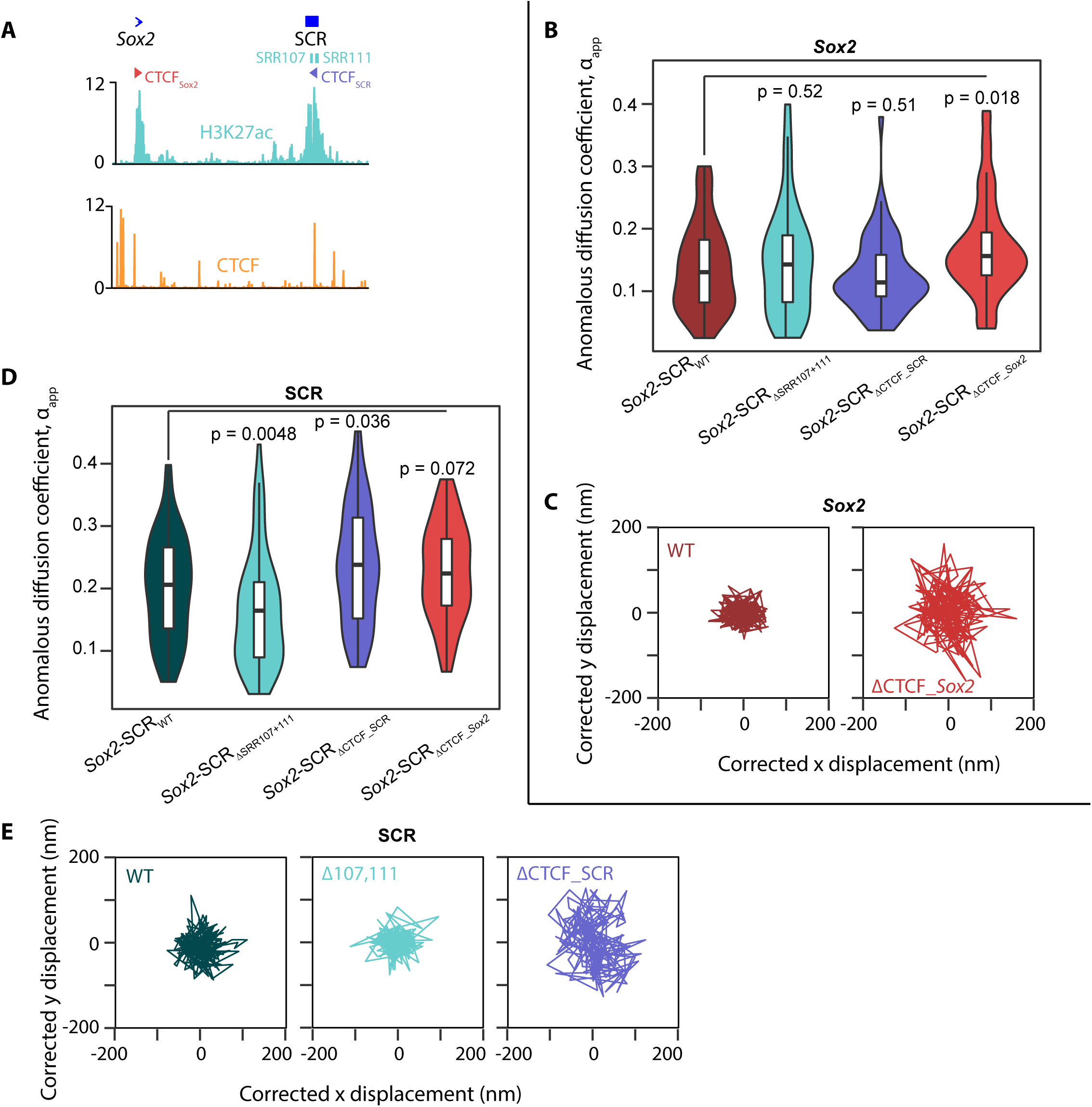
Loop extrusion and enhancer activity have opposing effects on chromatin dynamics. **a**) Overview of the *Sox2* locus, showing positions of the gene and SCR, and showing the regions deleted in the *Sox2-SCR_ΔSRR107+111_*, *Sox2-SCR_ΔCTCF_Sox2_*and *Sox2-SCR_ΔCTCF_SCR_* (shown in cyan, red and blue, respectively). Below are shown ESC ChIP-seq profiles for H3K27ac (cyan) and CTCF (orange). **b**) Violin plot showing distributions of apparent anomalous exponents measured from the individual movies for the *Sox2* promoter, comparing *Sox2-SCR_WT_*, *Sox2-SCR_ΔSRR107+111_*, *Sox2-SCR_ΔCTCF_Sox2_*and *Sox2-SCR_ΔCTCF_SCR_* ESCs. Comparisons are made by Wilcoxon rank sum tests, with *p*-values given. **c**) Sample tracks of the 2D trajectories for *Sox2* in *Sox2-SCR_WT_* and *Sox2-SCR_ΔCTCF_Sox2_* cells, plotted on the same scale. Displacements have been corrected for substrate displacement, and the plots have been centered on the median positions. **d**) Violin plot showing distributions of apparent anomalous exponents measured from the individual movies for the SCR, comparing *Sox2-SCR_WT_*, *Sox2-SCR_ΔSRR107+111_*, *Sox2-SCR_ΔCTCF_Sox2_*and *Sox2-SCR_ΔCTCF_SCR_* ESCs. Comparisons are made by Wilcoxon rank sum tests, with *p*-values given. **e**) Sample tracks of the 2D trajectories for the SCR in *Sox2-SCR_WT_*, *Sox2-SCR_ΔSRR107+111_* and *SCR_ΔCTCF_SCR_* cells, plotted on the same scale. Displacements have been corrected for substrate displacement, and the plots have been centered on the median positions.

To assess this possibility further, we ran molecular dynamics simulations of the above genetic perturbations within our framework, by either converting one or other of the CTCF-bound beads to a “neutral” one to mimic specific loss of the binding motif, or recapitulating the *Sox2-SCR_ΔSRR107,111_*line by deleting the corresponding beads, and abolishing PolII binding at the inactivated *Sox2* promoter. The original model did not recapitulate the chromatin dynamics changes observed in the experiments (**Fig S8**). In particular, the model predicted that transcription factor site deletions at the SCR would increase mobility at the enhancer, instead of placing greater constraints as we had observed. Strikingly, better agreement to experimental observations was accomplished by increasing RNA polymerase stability at bound enhancers, effectively reinforcing the competition between transcription-linked compartmentation and loop extrusion (see Methods). In these simulations, removal of the local transcription factor binding sites placed greater constraints on the enhancer, and removal of CTCF had the inverse effect (**Fig 7a,b**). Moreover, these simulations predicted greater cohesin occupancy at the SCR CTCF site on deletion of the flanking transcription factor binding sites. This may be predicted to allow more loop extrusion to pass through the locus, resulting in greater average local constraints as more alleles have stalled cohesin at the adjacent CTCF site. In support, we found by allele-specific ChIP-qPCR that binding of the cohesin subunit Rad21 is increased at the SCR CTCF site when adjacent transcription factor binding regions, SRR107 and SRR111, are deleted in *cis* (*p* = 0.001; two-tailed t-test; **Fig 7c**), suggesting that these elements are indeed antagonistic to cohesin recruitment and/or loop extrusion. Overall, these results extend on recent studies into the effects of loop extrusion and its stalling by CTCF on chromatin dynamics, showing a surprising localized effect. Furthermore, two different mechanisms believed to contribute in building the same promoter-enhancer topology, CTCF-mediated architectures and enhancer-promoter “contacts”, presumably via specific transcription factors and potential compartmentation, may actually have opposing effects on local chromatin dynamics (**Fig 7d**).

**Figure 7.**
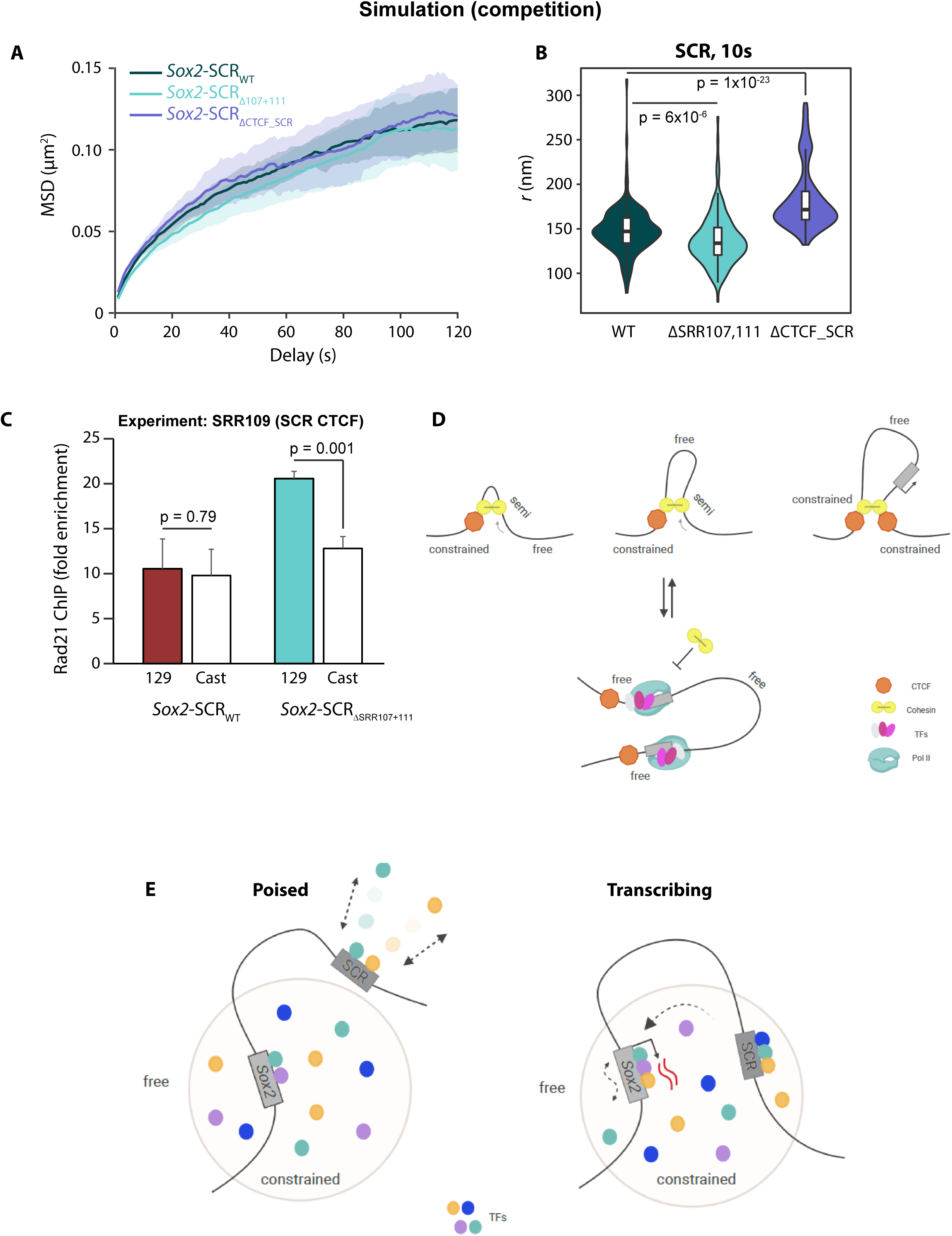
Competition between loop extrusion and transcriptional compartmentation can explain locus dynamics. **a**) Ensemble MSD curves for the SCR, derived from molecular dynamics simulations of the *Sox2* locus in wild-type or SCR mutant conditions, after factoring in reinforced competition between cohesin-mediated loop extrusion and RNA polymerase II clustering. Lines show median values and shading indicating the median absolute deviation. **b**) Violin plots showing distributions of apparent explored radii of the SCR after 10 s, derived from the same simulations as **a**. Reduction in mobility of *Sox2-SCR_ΔSRR107+111_* cells, and increased mobility in *SCR_ΔCTCF_SCR_* cells, compared to wild-type, is assessed by Wilcoxon rank sum tests, with *p*-values given. **c**) ChIP-qPCR quantification, expressed as fold enrichment over a negative control region, of amount of Rad21 binding at the musculus/129 (colored) or castaneus (white) alleles of the SCR CTCF site in *Sox2-SCR_WT_* and *Sox2-SCR_ΔSRR107+111_*cells. Comparisons between allelic binding are made by two-tailed t-tests, with *p*-values given. More Rad21 is bound to the SCR CTCF site, specifically on the allele where SRR107 and SRR111 are deleted. **d**) Schematic of competing processes dictating dynamics at the SCR. At the top, extruding cohesin complexes (yellow) place temporary local constraints on mobility of the bound sequences. Such constraints are stabilized when the cohesin encounters bound CTCF (orange) in the appropriate orientation. At the bottom, bound RNA polymerase and transcription factors can also coordinate the *Sox2* and SCR position, but are refractory to cohesin recruitment and/or extrusion, hence maintaining a relatively more mobile chromatin locus. The equilibrium of these two processes, shifted by deletion of the flanking transcription factor binding sites in *Sox2-SCR_ΔSRR107+111_*cells, results in more constrained chromatin on average. **e**) Two-step model of *Sox2* transcription, and its consequences for locus mobility. In the poised state (left), the *Sox2* gene resides within a nuclear hub of concentrated transcription factors, whereas the SCR is outside and exchanging with less concentrated factors within the nucleoplasm. Due to its dense environment, the hub places greater constraints on local chromatin mobility. In the transcribing state (right), the enhancer is more constrained on entering the nuclear hub, and this more efficient exchange with the gene allows transcriptional firing. The gene, while remaining constrained within the hub, has more local kinetic energy as a potential consequence of transcription itself. Note that the physical distance between SCR and *Sox2* is less important in this model than whether or not the elements are residing in the nuclear hub.

## DISCUSSION

By combining live imaging of labeled endogenous loci with a battery of perturbation experiments and molecular dynamics simulations, we have teased apart some of the spatiotemporal complexities of enhancer-promoter communication. The ANCHOR system employs relatively small tags that can be placed extremely close to genomic loci of interest with negligible observable effects on transcription, nuclear organization and cell state, so is a valuable tool for such studies. The first question we addressed was whether promoters and enhancers need to directly juxtapose for transcriptional firing, since conventional C-method and light microscopy approaches have given rather different viewpoints^20^. We found that *Sox2* and the SCR are frequently in close proximity (distances < 200 nm), and significantly closer than control regions of identical genomic separation, in agreement with Hi-C predictions. However, transcriptionally active loci can be also be found with greater physical separation between the *Sox2* and SCR than non-transcribing alleles, strongly arguing against any obligation for their direct juxtaposition for expression. Moreover, promoter-enhancer distances were unchanged in all transcriptional perturbations in this study that did not affect the TAD border at the SCR, completely in line with recent C-method studies at this locus^44, 46^. So if direct juxtaposition of the enhancer is not required for transcription, how close is close enough? Clusters of PolII and transcription factors have been observed to have diameters of ∼100-200 nm in live nuclei^12, 13^, providing a reasonable first estimate of the size of proposed transcriptional hubs, and thus the maximum separation of regulatory elements if they need to share the same microenvironment for transcriptional firing. What is unclear so far is whether the enhancer needs to be engaged in the hub during the whole transcriptional cycle, or if it is free to disengage after initiation in a “hit and run” mechanism. Since control sequences separated by only 150 kb are also frequently closer than the proposed 200 nm threshold, and these measurements are below the diffraction limit of our microscopy setup, we could not resolve these models in this study. However, since the SCR is less constrained than *Sox2* in non-transcribing alleles, but has average dynamics indistinguishable from the promoter when the gene is transcribed, our results indirectly favor enhancer engagement in the transcriptional hub/environment on at least the minute scale, in line with studies in a Drosophila transgene^7^. Studies in loci with more distal enhancers, particularly those which seem to depend on cohesin-mediated loop extrusion for gene activation (presumably as a mechanism to help the enhancer “find” its distal target)^42, 43^, in combination with visualization of any hubs themselves, will be required to formally address these questions.

Whereas average inter-probe distances only demonstrated subtle differences at best in this study, MSD plots revealed more dramatic locus-and condition-specific differences in chromatin dynamics, with the most obvious being the greater relative freedom of movement of control sequences compared to the constrained promoter and enhancer. Conventional fits to ensemble log-transformed MSD curves are not precise and only robustly identify very flagrant differences in dynamics; our previously developed GP- FBM framework demonstrably gives greater precision in diffusive property measurements of single trajectories, both on simulations^25^ and in complex and noisy experimental data (see fits in **Fig S5a**). A limitation of this approach, which is also shared by radial MSD analysis, is the requirement to use cross-correlations of co-measured trajectories to correct for global nuclear movements and rotations, which can be quite large in ESCs (trials of alternative means to measure substrate movement revealed them to be much less robust). Since the movements of monomers are more highly correlated when they are closer along the polymer, the dynamics of loci separated by 150 kb are underestimated, resulting in anomalous diffusion exponents much smaller than expected from most polymer physics models^40^, also explaining the consistently lower measured *α_app_* values, when compared to results of our simulations, which do not have nuclear movements. These underestimated *α_app_* values are nevertheless precise and comparable between experimental conditions, allowing quantitative comparisons of chromatin mobility changes. There is the possibility that a perturbation could affect the coupling of the two labeled loci more than the individual particle dynamics in themselves, which could be missed by GPTool, but this does not seem to be the case, since most perturbations were found to have localized effects on only SCR or *Sox2*, and not both together.

The major question we wished to address was how chromatin mobility is affected by transcription. For the *Sox2* locus, we found clear links between chromatin dynamics and gene expression: the constrained promoter, but not the SCR, was sensitive to strong transcriptional inhibition by either enhancer element deletion or triptolide treatment, becoming even less mobile. Furthermore, when directly comparing transcribing and non-transcribing alleles within minimally perturbed cells, *Sox2* promoter dynamics were relatively unaltered, whereas the SCR became more constrained upon transcriptional firing of its partner gene. A model that may explain these findings are that there are (at least) two distinct processes linking transcriptional control and local chromatin mobility: entry into a transcriptional hub, and transcription itself (**Fig 7e**). Due to apparent confinement within a more limited volume, delimited by the hub, or perhaps just because of the even more crowded environment, presence of a locus in a transcriptional hub would be expected to reduce its underlying chromatin mobility. Conversely, as proposed previously^30^, the ATP hydrolysis and/or PolII tracking associated with transcription may be expected to add kinetic energy to the local chromatin and thus increase its mobility. In this manner, the results can then be explained by the *Sox2* gene, a key pluripotency gene with many constitutive and tissue-specific transcription factor binding sites at its promoter and proximal enhancers, as frequently nucleating its own hub in ESCs, partly explaining its consistent low mobility. The SCR, in contrast, has dynamics behavior consistent with it specifically entering the hub (and hence increasing its constraint) on transcriptional activation. Although the SCR is itself transcribed^57^, this is at much lower levels than *Sox2*, so may not feature significantly in the measurements. Moreover, it is not clear if the SCR eRNA is co-transcribed with *Sox2*, so may not necessarily occur when the SCR is sharing a transcriptional hub with *Sox2*. While explaining our results, and coherent with results from different prior studies, this hypothesis requires experiments beyond current technological capabilities to formally test, simultaneously tagging hub components, nascent RNA and the promoter and enhancer over sufficiently long time periods to cover multiple transcription cycles while maintaining high temporal resolution. However, the results of our study alone are highly evocative in suggesting that some loci are able to nucleate their own transcription-competent microenvironments, whereas others need to “find” them, raising questions about how such a “hierarchy” is formed, and the mechanisms influencing the search time to access pre-existing hubs.

As *Sox2* and the SCR are strong TAD borders in ESCs, and cohesin-mediated loop extrusion has recently been shown to affect chromatin mobility^40^, much of their local constraints could perhaps be explained by mechanisms independent of transcription. Indeed, abrogation of CTCF binding caused localized increases in chromatin mobility, which was particularly noteworthy at the SCR, considering that deletion of this site had negligible changes on global chromatin conformation (as determined by population-average 4C-seq) or *Sox2* expression. We also note that identifying the localized changes in chromatin dynamics is only possible by the analytical framework that we have developed. Even more striking was the finding that deletion of transcription factor binding sites had opposing effects on local dynamics to CTCF site deletion. Assuming that transcription factor clustering is one aspect of chromosome compartmentation^50^, this is reminiscent of the known antagonism between loop extrusion and compartments in defining chromosome conformations (**Fig 7d**). Indeed, our molecular dynamics simulations required direct competition between the loop extrusion and transcription-linked interaction processes to recapitulate the experimental findings. Since transcription hubs are assumed to place greater movement constraints on the chromatin within, this model implies a) that stalled cohesin is quantitatively more constraining than residing within a hub; b) that transcription factor clusters impede loop extrusion without causing stalled cohesin. This latter point is supported by the recent finding that transcription start sites form cohesin-independent TAD borders^65^, although the exact mechanisms are unclear, particularly as SMC complexes are able to bypass RNA polymerase and other large chromatin complexes *in vitro*^66^. Whatever the exact mechanisms, the implications for transcriptional regulation are very important: loop extrusion is believed necessary to allow the more distal enhancers to search effectively for their targets, but may actually be a barrier itself to forming the transcriptionally competent microenvironment. More detailed studies on the interplay of these two processes in living cells will be highly challenging but fascinating.

The *Sox2* locus, containing an ESC-dedicated (super)enhancer ∼150 kb from the promoter, comprising clusters of binding sites for tissue-specific transcription factors oriented around one major CTCF site facing the gene, and with no other confounding genes within the locus, was a convenient model for the studies presented here. Although many of the organizational principles uncovered here likely apply to many genes, more systematic studies will be required to formally test this. For example, the limited deletion studies performed so far have already shown large differences in robustness of TAD structures to loss of specific CTCF sites^67^, which could also play out in their dynamics. The ANCHOR system and its derivatives, coupled to orthogonal systems to track nuclear hubs and loop extrusion processes, will open new avenues to understanding the spatiotemporal regulation of transcription.

## METHODS

### Cell lines and culture

Mouse F1 ESCs (*M. musculus*^129^ × *M. castaneus* female cells obtained from Barbara Panning) were cultured on 0.1% gelatin-coated plates in ES medium (DMEM containing 15% FBS, 0.1 mM MEM nonessential amino-acids, 1 mM sodium pyruvate, 2 mM GlutaMAX, 0.1 mM 2-mercaptoethanol, 1000 U/mL LIF, 3 µM CHIR99021 [GSK3β inhibitor], and 1 µM PD0325901 [MEK inhibitor]), which maintains ESCs in a pluripotent state in the absence of a feeder layer^68, 69^. The *Sox2-SCR_WT_*and *Inter-Down* lines were generated by allele-specific CRISPR/Cas9-mediated knock-in of ANCH1 sequence^51^ into the SCR or downstream control region, and ANCH3 sequence^26^ into the *Sox2* promoter-proximal region or intervening control region, respectively. Flanking homology arms were first introduced by PCR amplification and Gibson assembly into vectors containing the ANCH1 or ANCH3 sequence, and 1 µg of each vector were co-transfected with 3 µg of plasmid containing Cas9-GFP, a puromycin resistance marker and a scaffold to encode the two gRNAs specific to the two insertion sites (generated by the IGBMC molecular biology platform) in 1 million F1 ESCs with Lipofectamine-2000. SNPs remove the NGG PAM sequence at the *castaneus* allele, assuring allele-specific knock-in (sequences given in **Table S3**). Two days after transfection, the cells were cultured for 24 h with 3 µg/mL, then 48 h with 1 µg/mL puromycin to enrich for transfected cells, before sorting individual GFP-positive cells into 96-well plates and amplification of individual clones. Heterozygous clones with the correct sequences inserted in the *musculus* allele were screened by PCR and sequencing.

*Sox2-SCR_ΔSRR107+111_* cells were derived from the *Sox2-SCR_WT_* line, essentially as in ref 44. Briefly, *musculus*-specific deletion of SRR111 was first achieved by co-transfection with plasmids containing four allele-specific gRNAs and Cas9D10A-GFP using Lipofectamine-2000, then PCR screening as in ref 70. Heterozygous deletion clones were subsequently co-transfected with vectors containing two allele-specific gRNAs and Cas9-GFP to delete SRR107 (sequences given in **Table S3**).

*Sox2-SCR_ΔCTCF_Sox2_* and *Sox2-SCR_ΔCTCF_SCR_*cells were derived from the *Sox2-SCR_WT_* line, essentially as in ref 61. 1 million cells were co-transfected with 2 µg plasmid containing Cas9-GFP, a puromycin resistance marker and a scaffold to encode a gRNA targeting the relevant CTCF site (generated by the IGBMC molecular biology platform; sequences given in **Table S3**) and 1 µg Alt-R homology-directed repair single-stranded DNA oligonucleotide (IDT; sequences given in **Table S4**) with Lipofectamine-2000. The repair oligonucleotides are designed to replace the CTCF motif with an *EcoR*I site to help screening knock-outs. Two days after transfection, the cells were cultured for 24 h with 3 µg/mL, then 48 h with 1 µg/mL puromycin to enrich for transfected cells, before sorting individual GFP-positive cells into 96-well plates and amplification of individual clones. Deletions were evaluated on agarose gel after PCR amplification of the region around the deletion and *EcoR*I digestion. Homozygous deletions were confirmed by sequencing.

*Sox2-SCR_MS2_* cells were derived from the *Sox2-SCR_WT_*line by co-transfecting 1 million cells with 2 µg plasmid containing Cas9 and the scaffold encoding a gRNA at the *Sox2* 3’UTR, and 2 µg targeting vector containing the MS2 cassette, comprising 24 copies of the v6 MS2 repeat^58^, a hygromycin resistance cassette flanked by loxP sites, and primer binding sites to facilitate screening for inserted clones, following the strategy of ref 71 (**Fig S9**), with Lipofectamine-2000. Three days after transfection, cells with integrated vector were selected with nine days of treatment with 200 µg/mL hygromycin. Individual cells were sorted into 96-well plates for clonal amplification, which were screened for heterozygous (*musculus*-specific) incorporation of full length 24xMS2 repeats by PCR and sequencing. To remove the selection marker, which may perturb *Sox2* mRNA function, 1 million cells were transfected with 4 µg Cre-GFP vector with Lipofectamine-2000, and after three days were subjected to selection with 6 µM ganciclovir for ten days. Marker excision and maintenance of full-length MS2 construct were verified by PCR and sequencing in surviving colonies. Finally, 1 million cells were co-transfected with 1 µg ePiggyBac transposase expression plasmid (System Biosciences) and 1 µg each epB-MCP-mScarletI, epB-OR1-EGFP and epB-OR3-IRFP vectors (the OR vectors are derived from those used in ref 25 by adding flanking inverted terminal repeats for recognition by ePiggyBac transposase; original vectors available from NeoVirTech) with Lipofectamine-2000. Nine days after transfection, fluorescent cells were sorted in bins (low, medium and high) based on expression of the different fluorescent proteins, and tested by microscopy. Cells with Scarlet^high^/GFP^low^/IRFP^low^ fluorescence were found to be optimal for imaging experiments, and this population was maintained for subsequent experiments.

### Live imaging

Cells were transiently transfected with OR vectors (except for the stably expressing *Sox2-SCR_MS2_* line) and imaged essentially as in ref 25. 150,000 cells were plated two days prior to imaging onto laminin-511-coated 35 mm glass bottom petri dishes and transfected with 2 µg each OR1-EFGP and OR3-IRFP plasmids with Lipofectamine-2000. Medium was changed just before imaging to remove dead cells. Imaging was performed on an inverted Nikon Eclipse Ti microscope equipped with a PFS (perfect focus system), a Yokogawa CSU-X1 confocal spinning disk unit, two sCMOS Photometrics Prime 95B cameras for simultaneous dual acquisition to provide 95% quantum efficiency at 11 µm x 11 µm pixels and a Leica Objective HC PL APO 100x/1.4 oil. A Tokai Hit Stage incubator allowed for maintenance at 37°C and 5% CO_2_ throughout the experiment. The system was controlled using Metamorph 7.10 software. For double-label experiments, EGFP and IRFP were excited by 491-nm and 635-nm lasers. Green and far-red fluorescence were detected with an emission filter using a 525/50 nm and 708/75 nm detection window, respectively. Time-lapse was performed in 2D, acquiring 241 time points at 0.5 s interval. For triple-label experiments, EGFP and IRFP were first imaged as for the double-label, then mScarletI was excited at an identical z-position with a 561-nm laser, and red fluorescence was detected with an emission filter using a 609/54 nm detection window. Time-lapse was performed in 2D, acquiring 301 time points at 1 s interval. For assessment of transcriptional bursting of *Sox2-SCR_MS2_* cells, mScarletI was excited with a 561-nm laser, and red fluorescence was detected with an emission filter using a 609/54 nm detection window. Ten positions were taken per acquisition session, obtaining 20 z-stacks of 0.5 µm interval for each position every 2 min, for a total of 2 hr. All time lapse was performed with Perfect focus system (PFS) of the microscope to avoid any focus drift.

### ESC differentiation

ESCs were passaged onto laminin-511-coated 35 mm glass bottom petri dishes, transfected with OR vectors as previously, and switched to DMEM (4.5 g/L glucose) supplemented with 2 mM GlutaMAX, 10% FBS, 0.1 mM MEM non-essential amino acids, 0.1 mM 2-mercaptoethanol without LIF or 2i inhibitors. After 24 h, the medium was changed daily and supplemented with 5 µM retinoic acid.

### Triptolide treatment

Two days after transfection with OR vectors, as previously, cell medium was changed for fresh medium, supplemented with either 500 nM triptolide or 1:2000 dilution of DMSO and incubated for 4 h prior to image acquisition.

### qRT-PCR

Total RNA was extracted using the Nucleospin RNA extraction kit (Macherey-Nagel) and reverse-transcribed using random hexamer primers and SuperScript IV (Invitrogen). qRT-PCR was performed on 10 ng cDNA aliquots in technical triplicates per biological replicate (three minimum) using QuantitTect SYBR Green PCR kit (Qiagen). Amplification was normalized to *SDHA* (succinate dehydrogenase complex flavoprotein subunit A). SNPs allowed primer design to distinguish *musculus* and *castaneus Sox2* expression^44^. Primer sequences are given in **Table S5**. Statistical comparisons were made by two-tailed t-tests.

### Cell cycle analysis

1 million cells were fixed in 66% ethanol on ice for 2 h, washed with PBS, then resuspended in 750 µL 50 µg/mL propidium iodide and 2 µg/mL RNase A in PBS, before incubation at 37°C for 30 min. Analysis of propidium iodide staining with a FACS Fortesa (BD Biosciences) allowed cells to be gated to G1 (2n DNA content), S (2n < DNA content < 4n) and G2/M (4n DNA content) phases. Statistical comparisons were made by two-tailed t-tests.

### Allele-specific 4C-seq

4C-seq was essentially carried out as in ref 44. Cells were fixed with 2% paraformaldehyde in 10% FBS in PBS for 10 min at 23°C. The fixation was quenched with cold glycine at a final concentration of 125 mM, then cells were washed with PBS and permeabilized on ice for 1 h with 10 mM Tris-HCl, pH 8, 100 mM NaCl, 0.1% NP-40, and protease inhibitors. Nuclei were resuspended in *Dpn*II restriction buffer at 10 million nuclei/mL concentration, and 5 million nuclei aliquots were further permeabilized by treatment for 1 h with 0.4% SDS at 37°C, then incubating for a further 1 h with 3.33% Triton X-100 at 37°C. Nuclei were digested overnight with 1500 U *Dpn*II at 37°C, then washed twice by centrifuging and resuspending in T4 DNA ligase buffer. *In situ* ligation was performed in 400 μL T4 DNA ligase buffer with 20,000 U T4 DNA ligase overnight at 23°C. DNA was purified by reverse cross-linking with an overnight incubation at 65°C with proteinase K, followed by RNase A digestion, phenol/chloroform extraction, and isopropanol precipitation. The DNA was digested with 5 U/μg *Csp*6I at 37°C overnight, then re-purified by phenol/chloroform extraction and isopropanol precipitation. The DNA was then circularized by ligation with 200 U/μg T4 DNA ligase under dilute conditions (3 ng/μL DNA), and purified by phenol/chloroform extraction and isopropanol precipitation. For *musculus*-specific 4C profiles, samples of the DNA were digested with *Ava*II, cutting specifically at the region between the *Csp*6I site and the non-reading primer annealing site on the *castaneus* allele. 100 ng aliquots of treated DNA were then used as template for PCR with bait-specific primers containing Illumina adapter termini (primer sequences in **Table S6**). PCR reactions were pooled, primers removed by washing with 1.8× AMPure XP beads, then quantified on a Bioanalyzer (Agilent) before sequencing with a HiSeq 4000 (Illumina). Sequencing read fastq files were demultiplexed with sabre (https://github.com/najoshi/sabre) and aligned to the mm10 genome with Bowtie^72^, and intrachromosomal reads were assigned to *Dpn*II fragments by utility tools coming with the 4See package^54^. 4See was also used to visualize the 4C profiles. Interactions were called for each replicate with peakC^73^ with window size set to 21 fragments, and were then filtered to only include the regions called as interacting across all wild-type replicates. For statistical comparison of specific interactions, the 4C read counts within 1 Mb of the bait for all replicates and conditions (from the same bait) were quantile normalized using the limma package^74^. The means of summed normalized 4C scores over tested interacting regions were taken as “interaction scores”, and were compared across conditions by two-tailed t-tests.

### 3D DNA FISH

FISH probes were generated by nick translation from fosmids centered on the ANCHOR insertion sites (WI1-2125O9 for *Sox2*, WI1-788C1 for *Inter*, WI1-111C20 for *SCR*, WI1-156F5 for *Down*), labeling the DNA with dUTP conjugated to biotin (*SCR* and *Down* probes) or digoxigenin (*Sox2* and *Inter* probes). For each hybridization experiment, 300 ng each probe was combined with 3 µg mouse *C_0_t* DNA and 20 µg yeast tRNA and resuspended in 5 µL hybridization mix (10% (w/v) dextran sulphate, 50% formamide, 1% Tween-20 in 2x SSC). Probe was denatured at 95°C for 5 min just before application to cells. FISH was performed as in ref 75. Briefly, cells were plated on 0.1% gelatin-coated coverslips in ES medium and fixed after 5 h (before colony formation) for 10 min with 4% paraformaldehyde in PBS, then permeabilized for 20 min in 0.5% (w/v) saponin, and incubated for 15 min with 0.1 N HCl, with washes in PBS between each incubation. Cells were then washed in 2x SSC and equilibrated in 50% formamide/2x SSC, before co-denaturing nuclei and probes at 85°C for 5 min, then hybridizing overnight at 37°C in a humidified chamber. Cells were washed three times with 50% formamide/2x SSC at 45°C for 5 min each, then three times with 1x SSC at 60°C for 5 min each, before cooling to room temperature in 0.05% Tween-20/4x SSC and blocking for 20 min in 3% BSA in 0.05% Tween-20/4x SSC. Anti-digoxigenin-rhodamine and fluorescein-avidin DN were diluted 1:100 in the same blocking solution and incubated with the cells for 1 h. Cells were washed three times for 5 min in 0.05% Tween-20/4x SSC, then coverslips were mounted in DAPI-containing Vectashield mounting medium. Interphase nuclei were imaged on a LSM 880 AxioObserver (Olympus) microscope, using a x63/1.4 objective lens (C plan-Aprochromat 63x/1.4 Oil DIC UV-VIS-IT M27), under 2x zoom and line-averaging 4 settings. The interval between z-slices was 0.37 µm and the x- and y-pixel size was 132 nm. Nuclei were segmented and inter-probe 3D distances were measured by custom scripts in ImageJ^76^. Distance distribution differences were assessed by Wilcoxon rank sum tests.

### Single-molecule RNA FISH

MS2v6 and *Sox2* probes were designed with Stellaris Probe Designer and labeled with Cy3 and Cy5, respectively, on the 3’ ends. Glass coverslips were placed into a 12-well plate and coated overnight with Poly-D-lysine. 400,000 cells were seeded per well and allowed to attach for 3 h to 70-80% confluency at 37°C, 5% CO_2_. Cells were washed three times with pre-warmed HBSS buffer (Hanks’ Balanced Salt Solution; no calcium, magnesium or phenol red; Thermo Fisher), before aspiration of buffer and cell fixation in 4% paraformaldehyde in PBS for 10 min at 23°C, then two washes in PBS for 10 min at 23°C. Cells were permeabilized overnight in 70% ethanol at 4°C, then washed for 5 min at 37°C with pre-warmed wash buffer (10% (v/v) formamide in 2x SSC). Coverslips were aspirated and dried for 5 min, before applying hybridization mix (50 nM each probe in 10% (w/v) dextran sulphate sodium salt, 10% (v/v) formamide, 2x SSC) and incubating overnight at 37°C in a humidified chamber. Coverslips were washed three times in wash buffer for 30 min at 37°C, then rinsed in 2x SSC and washed in PBS for 5 min at 23°C. Samples were mounted in Prolong Gold with DAPI (Invitrogen) on glass slides, and left to dry in the dark for 24 h before imaging on a Zeiss AxioObserver 7 inverted wide-field fluorescence microscope with LED illumination (SpectrX, Lumencor) and sCMOS ORCA Flash 4.0 V3 717 (Hamamatsu). A 40x oil objective lens (NA 1.4) with 1.6x Optovar was used. 27 z-stacks with 300 nm slices and 1×1 binning were taken with an exposure time of 500 ms for Cy3, 750 ms for Cy5 (100% LED power for both), and 25 ms for DAPI (10% LED power). Micromanager 1.4 software was used. A custom Python script was used to detect, localize and classify the spots (https://github.com/Lenstralab/smFISH). Cells and nuclei were segmented using Otsu thresholding and watershedding. Spots were localized by fitting a 3D Gaussian mask after local background subtraction^77^ and counted per cell.

### Western blot

∼ 3 million cells were collected by trypsinization, washed with PBS and lysed in 100 µL RIPA buffer (50 mM Tris-HCl pH 8, 150 mM NaCl, 1% (v/v) NP-40, 0.5% (w/v) sodium deoxycholate, 0.1% (w/v) SDS, 1 mM EDTA) for 30 min at 4°C. Cell lysate was sonicated with a Covaris E220 Bioruptor in AFA microtubes to shear genomic DNA (peak incident power 175 W, duty factor 10%, 200 cycles per burst, 150 s), before addition of 100 U benzonase (Sigma) in 100 µL RIPA buffer and incubation for 30 min at 30°C. Lysate was cleared by centrifugation at 22,000 *g* for 20 min at 4°C and protein concentration was measured with a Bio-Rad protein assay. 30 µg protein extract was loaded onto a 15% Bis-Tris polyacrylamide gel for electrophoresis then transfer to a PVDF membrane using a Mini Trans Blot Cell (Bio-Rad). Membranes were blocked for 1 h at 23°C in 4% milk/PBST (0.1% Tween-20 in PBS), then incubated with primary antibody (anti-SOX2, Santa Cruz #sc-365823; anti-histone H3, H31HH.3EI- IGBMC) at 1:1000 dilution in 4% milk/PBST overnight at 4°C. Membranes were washed four times for 5 min each at 23°C in PBST, then incubated with 1:10,000 goat anti-mouse-alkaline phosphatase (IGBMC) in 4% milk/PBST for 1 h at 23°C, before washing four times for 5 min each at 23°C in PBST and developing with Pierce ECL Western blotting substrate (Thermo Fisher). Image acquisition was performed on ImageQuant 800 (Amersham).

### ChIP-qPCR

For CTCF ChIP, 40 million ESCs were harvested with TrypLE (Invitrogen) and fixed for 10 min with 1% formaldehyde, then quenched with 125 mM glycine for 5 min at 23°C then 15 min at 4°C. Cells were washed with 10% FBS/125 mM glycine in 1x PBS, then lysed in 275 µL lysis buffer (50 mM Tris-HCl pH 8, 10 mM EDTA, 1% SDS, 1x complete EDTA-free protease inhibitors) on ice for 20 min. 320 µL lysate was transferred to a microtube 500 AFA and sonicated with a Covaris E220 (175 V peak incidence power, 20% duty factor, 200 cycles per burst) for 25 min to shear chromatin to a 200-700 bp range. Cell debris was removed by centrifugation for 15 min at 20,000 *g*, 4°C. Equal volumes of protein A- and protein G-Dynabeads (Invitrogen) were mixed and prepared by washing in ChIP dilution buffer (16.7 mM Tris-HCl pH 8, 1.2 mM EDTA, 167 mM NaCl, 1% Triton X-100, 0.01% SDS, 1x complete EDTA-free protease inhibitors), then finally resuspending in an equivalent volume of ChIP dilution buffer. 100 µg chromatin aliquots were diluted eight-fold in ChIP dilution buffer and pre-cleared for 90 min with 30 µL prepared protein A/protein G-Dynabeads at 4°C on a rotating wheel. The pre-cleared chromatin was then incubated overnight at 4°C with 10 µL of either rabbit IgG (1 mg/mL, IGBMC facility) or rabbit anti-CTCF (07-729, Millipore). Immunoprecipitated chromatin was then adsorbed to 100 µL prepared protein A/protein G-Dynabeads for 3.5 h at 4°C on a rotating wheel. The beads were then washed twice for 5 min at 4°C with low salt wash buffer (20 mM Tris-HCl pH 8, 150 mM NaCl, 2 mM EDTA, 1% Triton X-100, 0.01% SDS), twice for 5 min at 4°C with high salt wash buffer (20 mM Tris-HCl pH 8, 500 mM NaCl, 2 mM EDTA, 1% Triton X-100, 0.01% SDS), twice for 5 min at 4°C with LiCl wash buffer (10 mM Tris-HCl pH 8, 250 mM LiCl, 1 mM EDTA, 1% NP-40, 1% sodium deoxycholate), then twice for 5 min at 4°C with TE buffer (10 mM Tris-HCl pH 8, 1 mM EDTA). The immunoprecipitated chromatin was eluted by two incubations with 250 µL elution buffer (100 mM NaHCO_3_, 1% SDS) for 15 min at 23°C. Eluate and input samples were decrosslinked by treatment overnight at 65°C, then DNA purified by RNase A digestion, proteinase K digestion, phenol/chloroform extraction and ethanol precipitation. qPCR was performed with QuantitTect SYBR Green PCR kit (Qiagen), using dilutions of the input material to generate the standard curve. Primer sequences are given in **Table S7**.

For Rad21 ChIP, 40 million ESCs were fixed for 45 min with 2 mM succinimidyl glurarate, then for 10 min with 1% formaldehyde at 23°C, before quenching with 125 mM glycine for 15 min at 23°C. Cell pellets were washed twice with ice-cold PBS, then resuspension and incubation in lysis buffer 1 (50 mM HEPES-KOH pH 7.5, 140 mM NaCl, 1 mM EDTA, 10% glycerol, 0.5% NP-40, 0.25% Triton X-100) for 10 min at 4°C. Lysates were centrifuged for 5 min at 2000 *g*, 4°C, before incubation for 10 min at 4°C in lysis buffer 2 (10 mM Tris-HCl pH 8, 200 mM NaCl, 1 mM EDTA, 0.5 mM EGTA). Lysates were again centrifuged for 5 min at 2000 *g*, 4°C, before resuspending in ice-cold lysis buffer 3 (10 mM Tris-HCl pH 8, 100 mM NaCl, 1 mM EDTA, 0.5 mM EGTA, 0.1% sodium deoxycholate, 0.5% N-laurylsarcosine).

Chromatin was sonicated in a probe sonicator at 20 A (15 s on/30 s off) for 2.5 min at 4°C. Triton X-100 was added to the sonicated lysate to precipitate any remaining debris, and the cleared lysate was incubated overnight with 5 µg anti-Rad21 antibody (Abcam ab217678) at 4°C with rotation. 80 µL protein A/G Dynabeads (Invitrogen) was added to the antibody-bound chromatin and incubated overnight at 4°C with rotation. The immunoprecipitates were washed six times at 23°C with RIPA buffer followed by a TBS buffer wash, then bound chromatin was eluted by incubation for 30 min at 65°C with elution buffer (50 mM Tris-HCl pH 8, 10 mM EDTA, 1% SDS), before addition of 4 µL 20 mg/mL proteinase K and reversal of crosslinks by incubating at 65°C overnight. DNA was purified with phenol/chloroform extraction and ethanol precipitation before qPCR quantification (primer sequences given in **Table S7**), using serial dilutions of pre-immunoprecipitated input material to generate the quantification standard curve. Quantification at the musculus and castaneus SRR109 (SCR CTCF site) was expressed as fold-enrichment over the amplification at a non-genic, negative control sequence, not bound by Rad21, CTCF or known pluripotency transcription factors.

### Localizing and tracking chromatin loci

Live imaging experiments were treated essentially as in ref 25. First, spot detection and tracking was performed with ICY^78^, before localization precision enhancement in GPTool^25^. This assumes that the spots have the shape of a 2D Gaussian function, as follows,

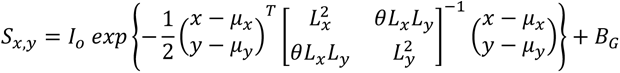

with *µ_i_* representing the center of mass of the spot, *L_i_* its size in directions x and y, -1 < *θ* < 1 a possible rotation, while *B_G_* and *I_o_* are background and spot signal, respectively. The localization is optimized using the NM-Simplex method, and localization error is estimated using the Metropolis-Hastings algorithm. Any outlier detection events, defined as having any of the signal intensity, spot size or localization error deviating from the median value within the experiment by more than twice the inter-quartile range, were removed from subsequent analysis. For the *Sox2-SCR_MS2_*line, ICY was additionally used for spot detection and tracking of MCP (mScarletI). The majority of experiments had MCP detection for virtually all or none of the frames, and were categorized as “transcribing” and “non-transcribing” cells, respectively. The other experiments were only included in the analysis if they contained at least 75 contiguous frames with MCP signal (“transcribing”) and/or absence (“non-transcribing”). For these, the largest contiguous stretch of frames above this threshold was maintained for subsequent analysis.

### Inter-probe distance analysis

For inter-spot distance measurements, effects of camera misalignments and chromatic aberrations were corrected with a set of generic affine transformations, including translation, rotation and scaling, defined as,

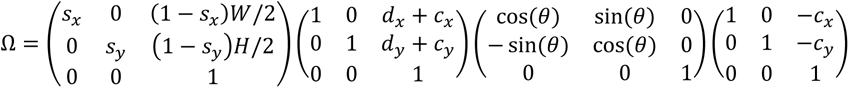

where *s_i_* accounts for scaling in directions x and y, *d_i_* accounts for translation in both directions and *θ* is the angle of rotation between both channels in relation to point *c_i_*. To infer optimal parameters for correction, five frames from each of the movies recorded in the session were used and the likelihood was maximized using the Nelder-Mead simplex method,

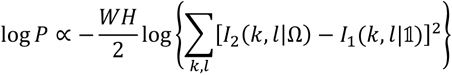

where *W* and *H* correspond to width and height of images, and *I_r_(k,l|A)* is the value of pixel (k,l) in channel r given transformation A. Here, #x1D7D9;#x1D7D9; represents the identity matrix. Visual inspection of acquired images before and after correction, and comparison with direct subtractions from optical beads, found the affine transformations to work better than the optical beads. After applying these transformations to the final localized spots, the 2D distances between two spots is then computed in a Pythagorean framework. The median distances were taken for each imaging experiment, and distributions of these distances were compared between cell lines and experimental conditions (or 2D distances determined by analogous methods such as FISH) by Wilcoxon rank sum tests.

### Transcriptional bursting measurements

Using the LiveCell pipeline (github.com/Lenstralab/livecell)^79^, maximal intensity projections were made of the 3D movies, followed by background subtraction and cell segmentation using Otsu thresholding and watershedding. Transcription bursts were detected and tracked using a 2D Gaussian fit followed by hysteresis filtering using 5 and 8 standard deviations of the background as low/high thresholds, respectively. Output data were manually checked to reject erroneous tracking or dividing cells.

### GP-FBM measurement of diffusive parameters

Diffusive parameters for individual trajectories were estimated using GPTool^25^, which assumes that the stochastic diffusion trajectory of a chromatin locus can be modeled as a Gaussian process with the following fractional Brownian kernel:

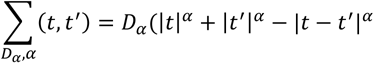

producing generalized Brownian motion with mean squared displacement,

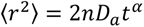

where *n* corresponds to the number of degrees of freedom. Formal derivation of the model and details of the methods used to optimize the model fit are found in ref 25 and references within. Experimental trajectories were verified as being reasonably fit by a fractional Brownian regime by checking that displacements are self-similar Gaussian distributed with the modeled covariance matrix (**Fig S3**) and that the velocity autocorrelation function from experimental trajectories,

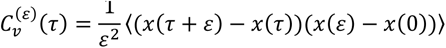

with velocity defined as *v(τ) = ε^-1^[x(τ+ε)-x(τ)]*, fits the theoretical velocity autocorrelation function for FBM (**Fig S4**):

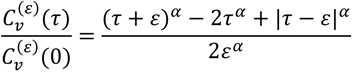

To measure and correct for substrate movement, GPTool builds a covariance model to handle the cross-correlation that substrate motion introduces into the two measured particles’ trajectories, generating the probability distribution:

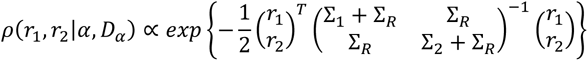

where *Σ_1_, Σ_2_* and *Σ_R_* are FBM covariance matrices for the two particles and substrate, respectively, with diffusion parameters *D_α_* = {D_α1_, D_α2_, D_αR_} and *α* = {α_1_, α_2_, α_R_}. Formal derivation of the model and details of the methods used to optimize the model fit are found in ref 25 and references within. Overall, GPTool outputs, for each individual imaging experiment: more precise trajectories for the two particles; estimations of diffusive parameters *D_α_* and *α* for each particle, with and without substrate correction; estimations of the trajectory of the substrate, as well as its diffusive parameters, D_αR_ and α_R_.

### Trajectory plots

To generate the individual trajectory plots (e.g. **Fig 3a**), the inferred substrate motion at each time interval was subtracted from the x and y coordinates of the GPTool-refined tracked particles, then multiplied by the scaling factor (110 nm per pixel) to obtain corrected coordinates in nm for the particle at each time point. The deviation of these values from the median x- or y-value, respectively, of the trajectory were plotted, maintaining their temporal order, to generate comparable trajectory plots across experiments centered on the origin.

### MSD analysis

MSD analysis was performed using the msdanalyzer utility in Matlab^80^. For each individual trajectory, the time-averaged MSD was computed (time steps of 0.5-120 s for most experiments; 1-300 s for experiments with the *Sox2-SCR_MS2_*line), using either the trajectories uncorrected or corrected by GPTool for substrate movement, as stated. Because some trajectories exhibited large deviations from the average, especially at long time, we chose to calculate the ensemble median of all individual MSD and the median absolute deviation as a measure of the variability. For testing the fit of GPTool-measured parameters to experimental data, theoretical 2D MSD curves were made from these measured parameters for each individual trajectory, giving either

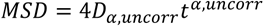

for parameters called without substrate correction, or

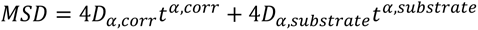

for the particle and substrate parameters called when substrate motion correction is applied. The ensemble medians of these MSD curves were plotted with the experimental MSD curves. Radial MSD was computed in exactly the same way as single-particle MSD, but using the evolution of Euclidean distance between two particles instead of the displacement of a single particle. The diffusive parameters were fit to the first 60 s of the log-log plots of radial MSD using linear least squares with bisquare weights, implemented in Matlab. 95% confidence intervals were estimated as two standard deviations of the point estimates. Diffusive parameter differences between *Sox2*-SCR and *Inter-Down* radial MSD were assessed by ANCOVA.

### Statistical comparison of inferred diffusive parameter distributions

The distributions of inferred apparent anomalous diffusion coefficients from different genomic loci or experimental conditions were compared by Wilcoxon rank sum test, with Benjamini-Hochberg multiple testing correction in cases where four or more statistical tests were combined. Since *Dα_app* values cannot be fairly compared across trajectories with different inferred values of *α_app_*, the inferred values of *D_α_app_* and *α_app_* for each particle trajectory were fed into the generalized Brownian motion equation to compute, for each individual trajectory, the expected radius of exploration, *r*, in a given time, *t*:

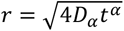

These were computed for all trajectories for different time-lags (1, 10, 30 and 60 s), and their distributions were statistically compared with Wilcoxon rank sum tests. Note that these use the substrate-corrected inferred diffusive parameters, so the resultant “expected” explored area is smaller than experimental observations.

### Molecular dynamics simulations

Molecular dynamics simulations were performed via the multipurpose EspressoMD package^81^, as in ref 56. Individual proteins are represented by “beads” interacting via phenomenological force fields and move according to the Langevin equation, and the chromatin fiber is represented as a chain of beads connected by bonds. The position of every bead in the system evolves according to the Langevin differential equation that encodes Newton’s laws in the case of thermal bath with the friction *γ* due to an implied solvent in the presence of forces between beads encoded by energy potential functions *U*^82, 83^. Langevin equations for all beads are simultaneously solved in EspressoMD using a standard Velocity-Verlet numerical algorithm. The potential connecting *i* and *i+1* beads of the fiber is a finitely extensible non-linear elastic (FENE) spring that adds up to a steric repulsion potential between non-adjacent sites of the polymer, the Weeks-Chandler-Andersen (WCA) potential:

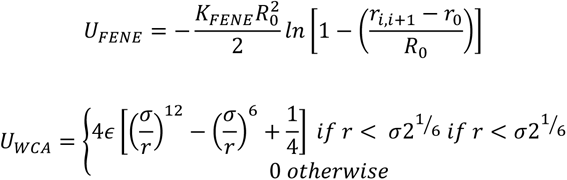

where *r_i,i+1_* is the distance between consecutive beads, and *σ* is where the interaction from repulsive becomes attractive and can be interpreted as the diameter of the particles. This value is a natural length scale of the system. In FENE the parameters are fixed to have an equilibrium distance of 1.6 *σ* with maximum extension of 0.8 *σ*, and a bond energy of *K_FENE_*= 30 k_B_T. Since the fiber is resolved at 1 kb, chromatin rigidity cannot be neglected (i.e. below the persistence length). Bending rigidity of the polymer is introduced via the Kratky-Porod potential for every three adjacent chromatin beads, where *θ* is the angle between three consecutive beads as given by:

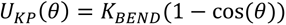

and *K_BEND_* is the bending energy. The persistence length in units of *σ* is given by *L_p_ = K_BEND_/k_B_T*. The model includes: (i) full 3D loop extrusion by interplay of cohesin dimers and CTCF, and (ii) transcription by PolII particles. To simulate association of cohesin and PolII with chromatin, a harmonic potential was employed, mimicking formation of a stable bond between two particles that fluctuate around an equilibrium distance *d_0_*:

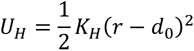

From now on, any description of the formation of a bond means the introduction of the aforementioned harmonic bond between the particles involved. To regulate the lifetime of these harmonic interactions, mechanisms of bond formation and removal were introduced according to cutoff distance *c_d_* below which a bond is formed, with a certain probability rate of detachment in units of time *τ_b_* = 2*τ* (*τ* being the fundamental unit of time in molecular dynamics; see below). These are then set to approximate the experimentally-observed range of PolII transcription and cohesin loop extrusion speeds and chromatin residence time. Introduction of the above mechanics is added on top of the strings-binders-switch model (SBS^84^), which encodes the association tendency of PolII with the *Sox2* promoter (defined as the 1 kb bead containing the transcription start site within the modeled region) by means of the shifted, truncated Lennard-Jones (LJ) potential, allowing spontaneous co-localization of beads around a distance *σ* with lifetime and stability properties depending on the depth of the energy well ε:

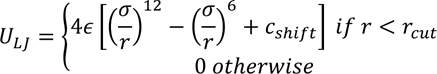

where *r_cut_* is a cutoff distance, *r* is the separation of any two beads, and *r_cut_* = 2.5*σ* for all LJ potentials in the simulations. This is a standard commonly used in the field to simulate phenomenological coarse-grained affinities^82, 83^.

For PolII interactions and transcription, the polymerase is presented as a bead with LJ interaction with specific beads of the chromatin fiber representing the promoter, enhancers (defined as beads containing H3K27ac peaks) or gene body (beads containing *Sox2* coding sequence) with energy ε = 2k_B_T. Such mild affinity helps to identify promoters as the correct sites where transcription initiation will take place (i.e. PolII forming stable bonds with the promoter bead) before the elongation process starts along the gene body. LJ interactions were also introduced among PolII beads (ε = 2.5k_B_T), to simulate their tendency to form condensates acting as transcription hubs, and between PolII and cohesin (ε = 2k_B_T) to simulate preferential loading of cohesin at promoter/enhancer beads. PolII transcription dynamics are simulated as a four-step process: attachment to the promoter in an exclusive manner, elongation starts, elongation proceeds through the gene body, detachment at the transcription termination site (the end of the gene body). A bond forms if the beads are less distant in space than a certain cutoff (2.7*σ*). To simulate the tendency of RNAPII to reel in gene body beads, a secondary bond is formed with the next bead on the chromatin fiber in the direction of transcription (*i*+1 bead, where *i* is the promoter coordinate on the fiber, and if transcription occurs in the sense direction; *i*-1 in the antisense direction). In the next step, RNAPII moves on the next site by forming new bonds with *i*+1 site and dissolving the old ones with *i*. This happens at a given rate (0.4 *τ_b_*^-1^) and only if the beads are found closer than a cutoff distance (*c_d_*= 1.05*σ*). These values are selected to obtain a RNAPII transcription speed approximately in the range of 1- 10 kbp/min observed experimentally. Upon reaching the TES, RNAPII stops and becomes unbound with rate 0.2 τb. Upon binding with promoters RNAPII loses its LJ interaction with promoters, since this is substituted by the bond itself. PolII is also allowed to form bonds with enhancers, although in this case no transcription procedure is initiated. As for PolII-promoter case, such bond dissolves at a given rate. Since cohesin and PolII, in their chromatin bound state, interfere with one another, we explored different rates of PolII-enhancer bond dissociation to study the effect of loop extrusion-transcription competition on the locus dynamics, ranging from 2 bonds lost per second (initial model, with limited competition) to 0.5 bonds lost per second (revised model, with reinforced competition). Cohesin concentration was maintained at 20 nM in the models.

For CTCF binding, a single bead interacts via LJ potential (ε = 2k_B_T) with specific beads of the chromatin fiber (CTCF ChIP-seq peaks containing oriented cognate binding motifs). Once a bond is formed (with rate 0.8 τb-1) it is pair-exclusive (i.e., other CTCF molecules cannot bind that same site). The bond dissolves at the rate 2×10^-5^ τb^-1^ and CTCF is again free to diffuse and search for other binding sites. Cohesin is represented as a pair of beads connected by one bond (*r_0_* = 1.6*σ* and *K* = 8k_B_T). Extrusion has three steps: attachment, active extrusion, and detachment. For attachment, each cohesin monomer forms a bond with the chromatin fiber. Bonds form when a cohesin monomer and a chromatin site come within a given cutoff distance (1.6*σ*) and at a certain rate (0.1 *τ_b_*^-1^). Only the case where both monomers simultaneously form bonds on adjacent chromatin beads is considered a successful attachment and the dimer is retained for the next step; otherwise bonds dissolve. If a promoter is already engaged in a bond with RNAPII, cohesin is forbidden to bind that promoter. Also, to favor cohesin loading in correspondence of active transcribing promoters, a 90% chance of binding has been introduced when a cohesin molecule is close to a promoter/enhancer and at the same time one RNAPII is close by as well (cutoff distance 1.5 σ), otherwise cohesin binding chance drops to 10%. The active extrusion and detachment steps follow the same mechanics as for RNAPII, with the difference that RNAPII can reel through cohesin bound sites while the inverse is not allowed. New bonds are formed if the distance is below 1.1σ. Such parameters produce ranges of cohesin extruding speed of 15-30 kb/min, which is within the range of experimentally observed values^47^. This allows cohesin to be affected by the surroundings during extrusion (e.g. presence of PolII close by or bound to adjacent sites). Cohesin detachment can occur independently at any step at a given rate (10^-4^ *τ_b_*^-1^) to fit its known chromatin residence time of 20 min. CTCF “loop anchors” are modeled so that cohesin cannot form new bonds with the next *i*±1 site if the latter is already bound by CTCF, provided it has the binding motif in convergent orientation. This renders extrusion dependent on CTCF dynamics. Finally, cohesin has LJ affinity for PolII both in the bound (ε = 2.5k_B_T) and unbound state (ε = 2k_B_T, higher affinity for bound RNAPII mimicking its suggested role in cohesin loading on chromatin^85^). PolII and loop extrusion dynamics are performed using a python script driving the EspressoMD library. The polymer initializes as a random walk and its dynamics first evolves in the absence of extrusion and transcription to generate an equilibrium coil formation. Extrusion and transcription processes are then switched on, and the evolution of the bead positions are tracked. Across all simulations, standard values for the friction coefficient (*γ* = 0.5) and the time step (t = 0.01) are used. To connect *in silico* space-time units with real distances and times, it is assumed that the concentration of DNA in the 3D simulation space is the same as that in a mammalian nucleus, giving a rough estimation of *σ* = 52 nm. For time units, the standard molecular dynamics relation *τ* = η(6πσ^3^/ε) is used. Assuming a viscosity ∼0.5P, the fundamental time unit is *τ* = 0.017 s. Concentrations of CTCF and PolII are taken from the physiological values and range from 30 to 60 nM. The energy scale of the system is given by the Boltzmann factor k_B_ multiplied by the temperature of the system T = 310 K.

The region modeled (mm10; chr3:34.3-35.1 Mb) encompasses the *Sox2* gene, 16 enhancer beads from the peaks of H3K27ac, and 6 CTCF binding regions with directionality determined by orientation of binding motifs. CTCF binding regions can be composed of up to 3 beads where CTCF occupancy signal is the highest (i.e. two regions upstream of *Sox2*), so giving a stronger anchoring function to such regions. Polymer models for *Sox2-SCR_ΔCTCF_Sox2_* and *Sox2-SCR_ΔCTCF_SCR_*have the relevant CTCF sites converted into non-CTCF binding beads, so to remove the cohesin anchoring effect. The model for *Sox2-SCR_ΔSRR107,111_*has the two enhancer beads immediately flanking the SCR CTCF region completely removed from the polymer, and reduced RNAPII stability to the promoter (from 0.02 bonds lost per second to 0.2 bonds lost per second), to mimic inactivation of the gene.

The positions of the beads corresponding to the positions of ANCHOR labels were tracked across 40 simulations of 1000 timepoints, and these trajectories were subsequently fed into MSD fits and GPTool diffusive parameter estimations, as for the experimental trajectories.

### Data sources

ESC and NPC Hi-C data were taken from ref 60, and re-analyzed and visualized using FAN-C^86^. The differential Hi-C map was derived as in ref 87. The normalized scores of the ESC Hi-C submatrix were subtracted from those of the NPC Hi-C submatrix, and the difference was expressed as a z-score: (diff – mean(diff))/standard deviation(diff)). Insulation was computed using FAN-C with a binsize of 7 bins (28 kb). ESC ChIP-seq datasets for H3K27ac and CTCF were obtained from the CODEX database^88^. For comparison of 2D *Sox2*-SCR distance distributions with analogous methods, results from oligoFISH were taken from ref 53, and results from CuO/TetO experiments were taken from ref 9.

## ACKNOWLEDGEMENTS

ANCHOR constructs were provided by NeoVirTech. The MS2v6 constructs were built by Biljana Atanasovska. Sequencing was performed by the Institute of Genetics and Molecular and Cellular Biology (IGBMC) GenomEast platform, a member of the France Génomique consortium (ANR-10-INBS-0009). This study was made possible because of the IGBMC Imaging Center (ici.igbmc.fr), flow cytometry and molecular biology platforms. BioRender was used for generating **Figs 7d** and **e**. This study was funded from grants by LabEx INRT (ANR-10-LABX-0030-INRT, a French State fund managed by the Agence Nationale de la Recherche under the frame program Investissements d’Avenir ANR-10-IDEX-0002-02), ERC (Starting Grant 678624 - CHROMTOPOLOGY) and Agence Nationale de Recherche (HUBDYN). AgP was supported by a fellowship from La Ligue Nationale Contre le Cancer. Work in the Mitchell laboratory was supported by the Canadian Institutes of Health Research (FRN PJT153186 and PJT180312), the National Institutes of Health (R01-HG010045-01), the Canada Foundation for Innovation, and the Ontario Ministry of Research and Innovation. Work in the Soutoglou laboratory was supported by the Academy of Medical Sciences (AMA). Work in the Lenstra lab was supported by an institutional grant of the Dutch Cancer Society and of the Dutch Ministry of Health, Welfare and Sport, and, Oncode Institute, which is partly financed by the Dutch Cancer Society. Work in the Papantonis lab was supported by the Deutsche Forschungsgemeinschaft via the SPP2191 and SPP2202 Priority Programs (Project Nos.: PA 2456/17-1 and PA 2456/11-2). Work in the Bystricky lab was supported by the Agence Nationale de Recherche and the Fondation ARC.

## AUTHOR CONTRIBUTIONS

AnP, AgP, KB and TS conceived and designed experiments. AnP and CE generated ANCHOR lines and performed nearly all experiments and particle tracking, with microscopy optimization assisted by EG. MB performed molecular dynamics simulations, with posterior analysis performed by BM. BM, GMO and NM performed most GPTool analyses. MACdK and TLL performed single-molecule RNA FISH and analysis, generated the core constructs for the *Sox2-SCR_MS2_* line and set up bursting analysis for AnP. KM and ES performed 3D DNA FISH, with analysis by SK. TT, VMS and JAM provided core constructs for the *Sox2-SCR_ΔSRR107,111_*line and performed allele-specific ChIP experiments. All authors contributed to writing of the manuscript.

## DECLARATION OF INTERESTS

The authors declare no competing interests.

## SUPPLEMENTARY INFORMATION

**Figure S1.**
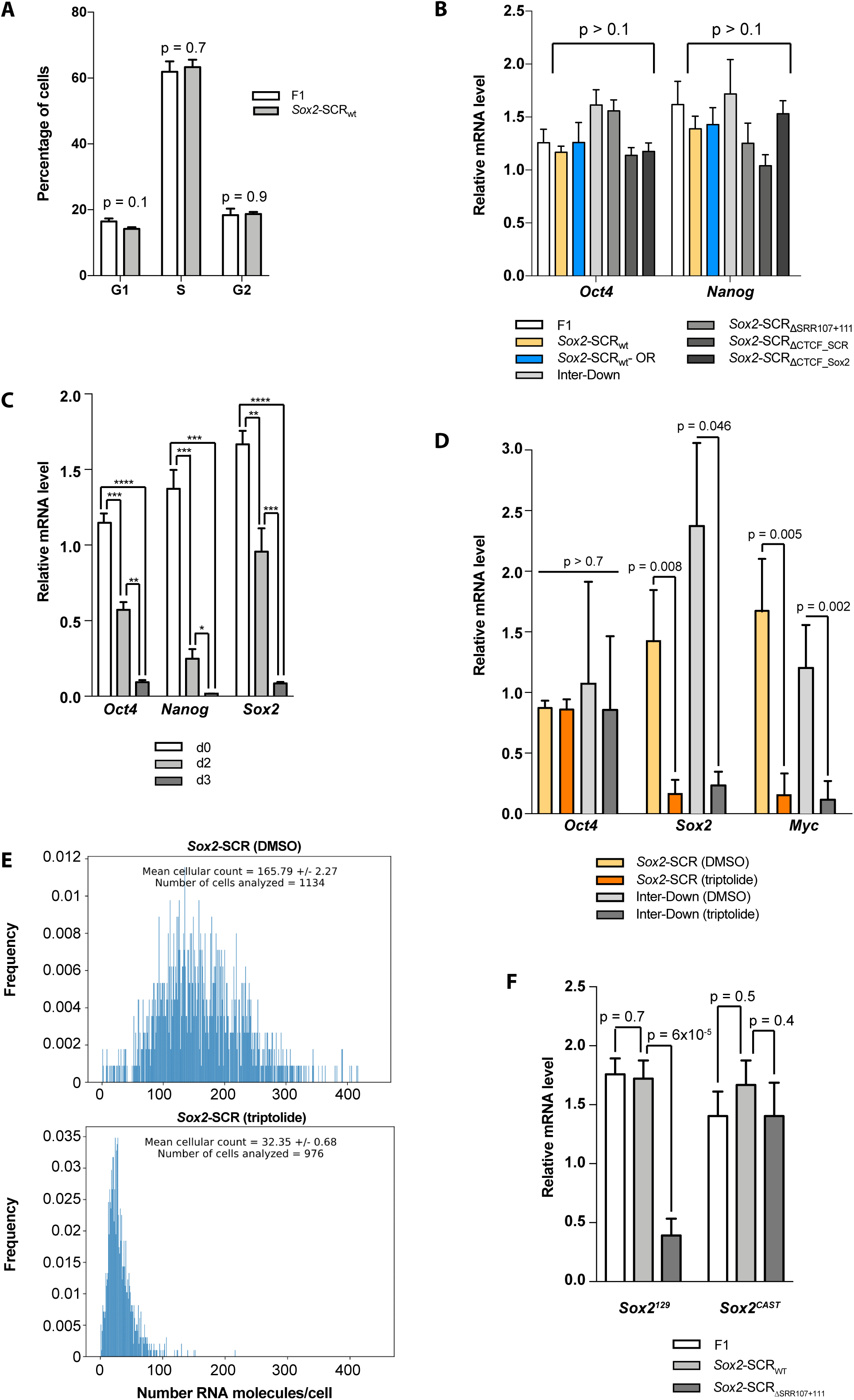
The ANCHOR system does not affect ESC function or response to transcriptional inhibition. **a**) Bar plot showing proportions of F1 and *Sox2-SCR_WT_* ESCs in G1, S and G2 phases of the cell cycle (*n* = 2). Error bars show standard deviations of the mean. Comparisons are made with two-tailed t-tests, with *p*-values given. **b**) qRT-PCR quantification of *Oct4* and *Nanog* expression, normalized to *SDHA*, in the different ESC lines used in this study, including *Sox2-SCR_WT_* cells with and without transfection of the ParB (OR) vectors (*n* >= 3). Error bars show standard deviations of the mean. Comparisons of expression with the F1 founder line are made by two-tailed t-tests, with minimum *p*- values given. **c**) qRT-PCR quantification of *Oct4*, *Nanog* and *Sox2* expression, normalized to *SDHA*, in *Sox2-SCR_WT_* cells after 0, 2 or 3 days of differentiation (*n* >= 3). Error bars show standard deviations of the mean. Comparisons are made by two-tailed t-tests (* *p* < 0.05; ** *p* < 0.01; *** *p* < 0.001; **** *p* < 0.0001). **d**) qRT-PCR quantification of *Oct4*, *Myc* and *Sox2* expression in *Sox2-SCR_WT_* and *Inter-Down* ESCs after acute treatment with DMSO carrier or triptolide, normalized to *SDHA* expression in DMSO-treated cells. Error bars show standard deviations of the mean. Paired comparisons between DMSO and triptolide treatments are made by two-tailed t-tests, with *p*-values given. *Oct4* mRNA has a longer half-life than the triptolide treatment time, unlike short-lived *Myc* and *Sox2* mRNA^89^. **e**) Histograms of single-molecule RNA FISH results for *Sox2* expression in *Sox2-SCR_WT_*cells treated with either DMSO or triptolide (>= 976 cells analyzed), showing numbers of mRNA molecules detected per cell. **f**) Allele-specific qRT-PCR quantification of *Sox2* expression, normalized to *SDHA*, for F1, *Sox2-SCR_WT_* and *Sox2-SCR_ΔSRR107+111_* cells, showing specific perturbation of *Sox2* transcription on the musculus allele where SRR107 and SRR111 are deleted. Error bars show standard deviations of the mean. Comparisons are made by two-tailed t-tests, with *p*-values given.

**Figure S2.**
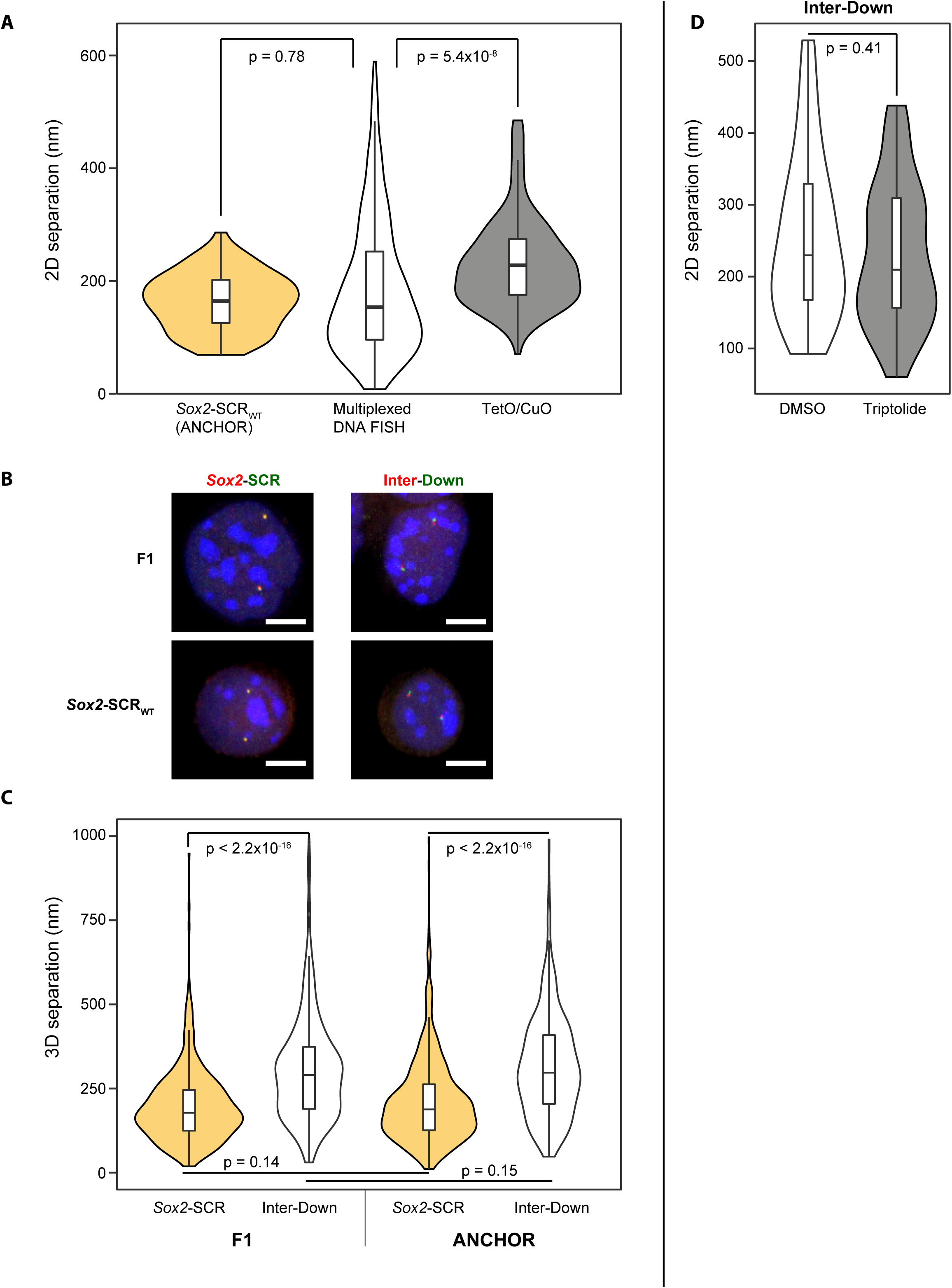
The ANCHOR system does not affect nuclear organization. **a**) Violin plots showing distributions of 2D *Sox2*-SCR distances, as measured by the ANCHOR system (this study), multiplexed DNA FISH^53^ and a TetO/CuO labeling system^9^. Comparisons are made by Wilcoxon rank sum tests, with *p*-values given. **b**) Representative maximal projection images of F1 or *Sox2-SCR_WT_* cells after 3D DNA FISH with probes for *Sox2* or Inter regions, labeled in red, and SCR or Down regions, labeled in green. DAPI DNA staining is shown in blue. Scale bar 5 µm. **c**) Violin plots showing distributions of 3D distances as measured by DNA FISH between *Sox2* and SCR, or between the Inter and Down control regions, in F1 and *Sox2-SCR_WT_* cells. Comparisons between distributions are made by Wilcoxon rank sum tests, with *p*-values given. Incorporation of *parS* sequences does not alter inter-probe distance distributions, and average Inter-Down distances are significantly greater than *Sox2*-SCR distances in both cell lines. **d**) Violin plots showing distributions of 2D distances between Inter and Down control regions in *Inter-Down* cells treated with either DMSO or triptolide. Comparison is made with Wilcoxon rank sum test (*p* = 0.41).

**Figure S3.**
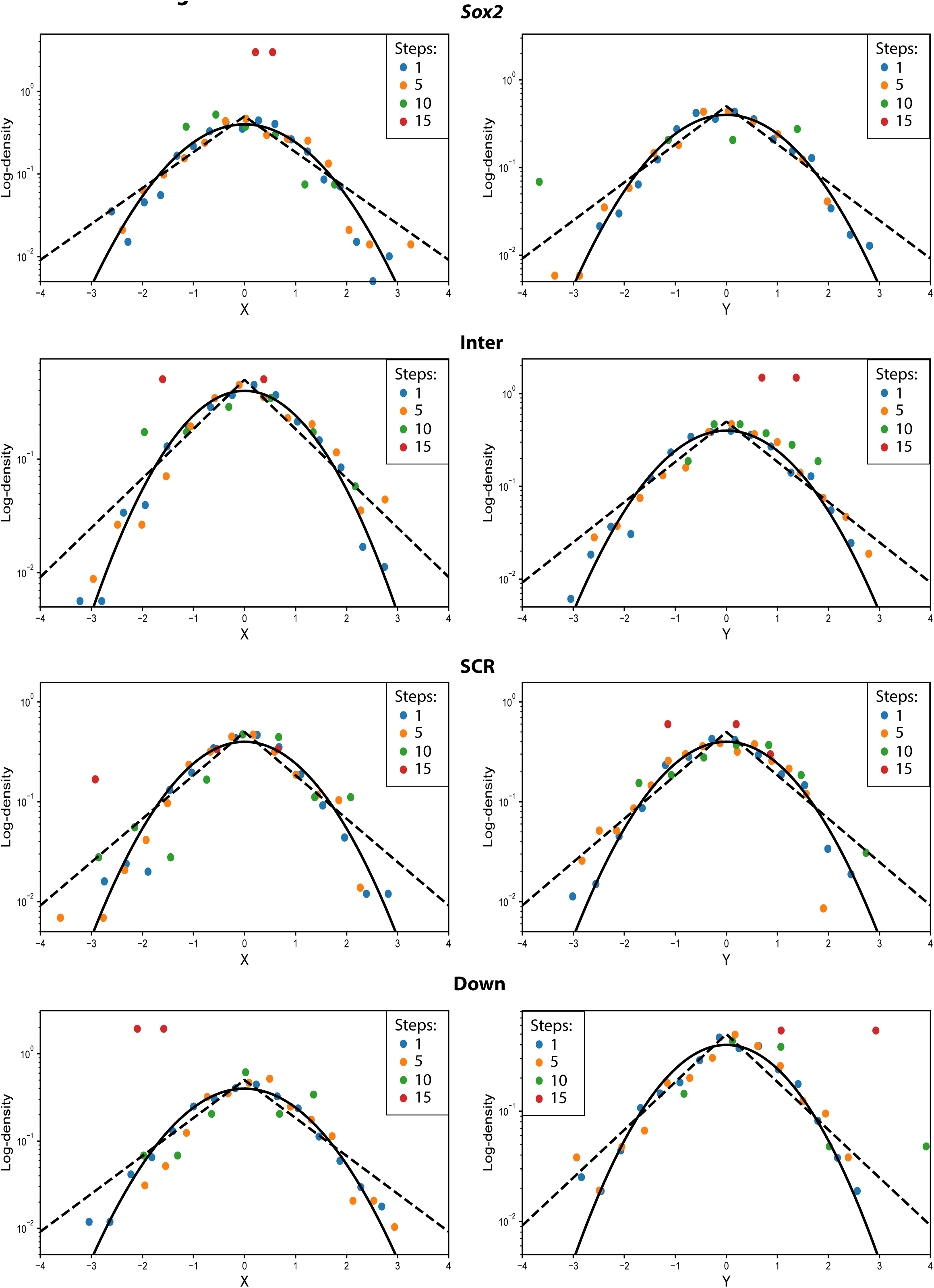
Gaussianity and self-similarity test on experimental trajectories. Displacements (left: x; right: y) were calculated and grouped for different time steps for the four labeled regions in *Sox2-SCR_WT_* and *Inter-Down* cells and compared against standard Gaussian (black line) and Laplacian (dashed line) distributions. All displacement values for a given time step *δ* were normalized by 2*D_α_δ^α^*, where *D_α_* and *α* were inferred from the corresponding trajectory. Experimental results show normally distributed and self-similar displacements over the time scales of our experiments, compatible with the FBM model.

**Figure S4.**
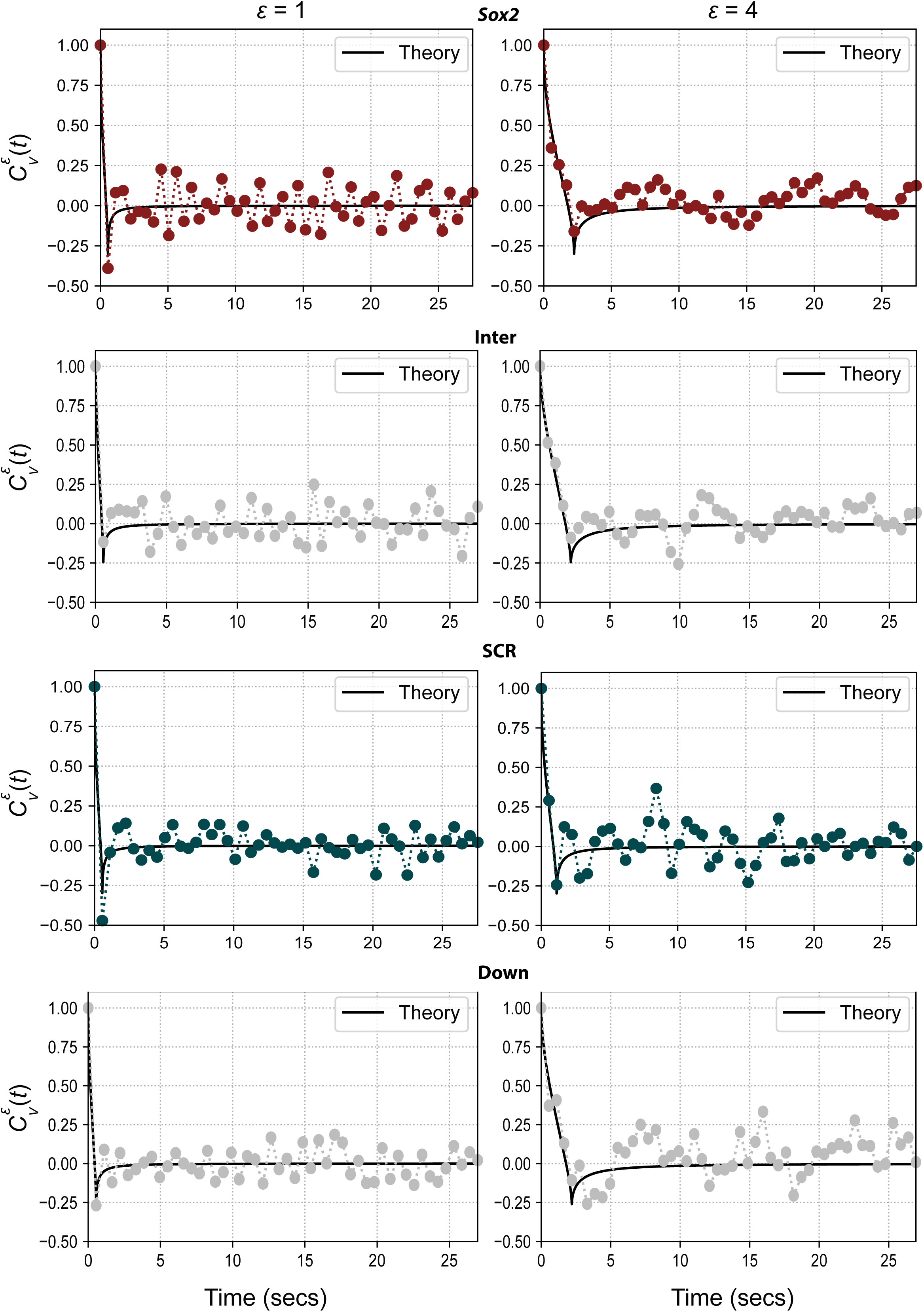
Experimental trajectories present FBM velocity autocorrelation. Velocity autocorrelation functions were calculated (see Methods) and grouped by labeled region in *Sox2-SCR_WT_*and *Inter-Down* cells to obtain smoother curves. Final results are compared to theoretical FBM curves (black line; see Methods) where the average anomalous exponent for each labeled region was used.

**Figure S5.**
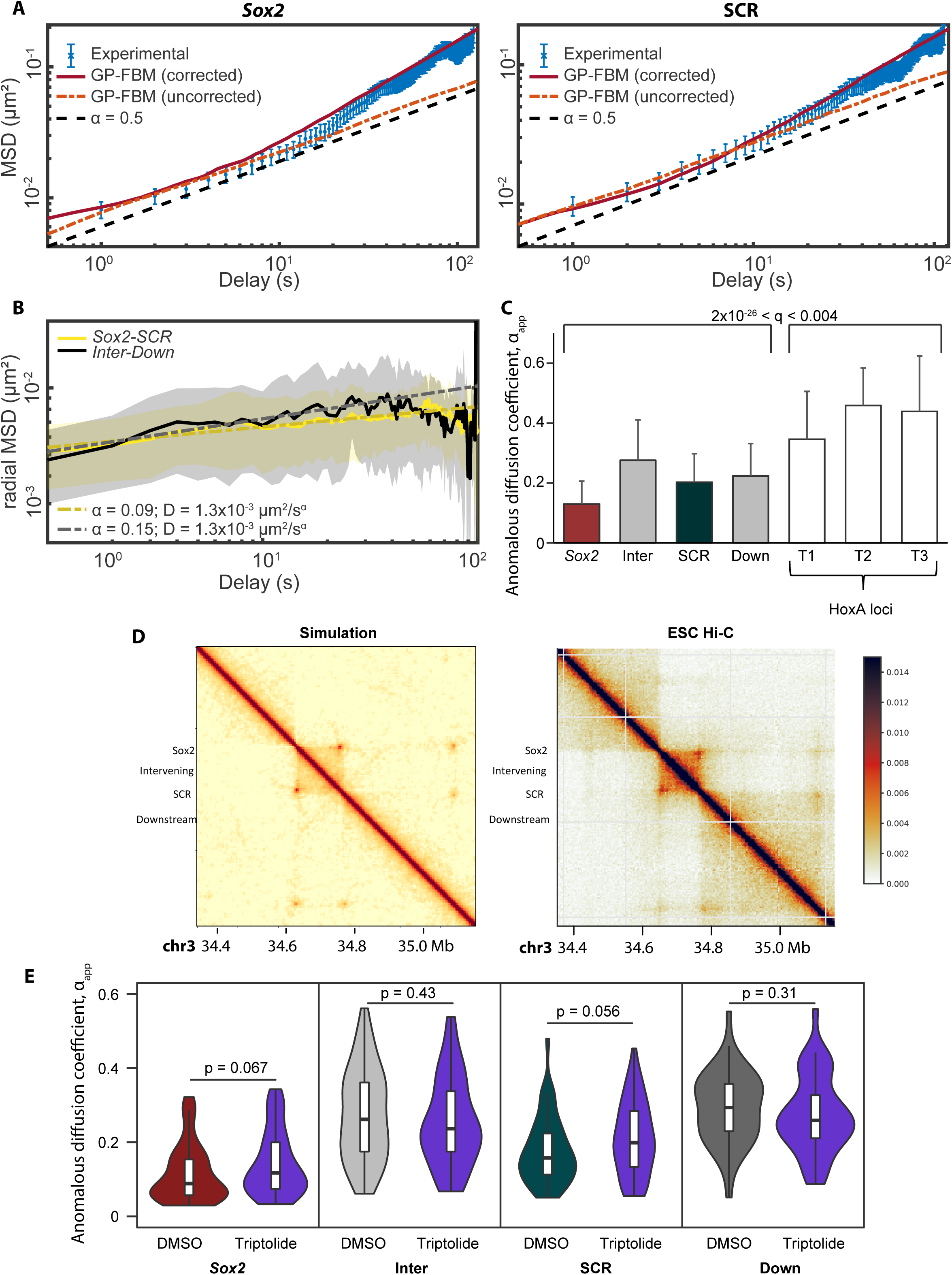
Substrate-corrected GP-FBM-measured diffusive parameters fit well to experimental trajectories. **a**) Uncorrected experimental median ensemble MSD curves (blue) are plotted for *Sox2* (left) and SCR (right), alongside median ensemble MSD curves derived from either fixing *α* at 0.5 (black), using the *D_α_app_* and *α_app_* parameters measured from GPTool without applying substrate correction (orange), or as the sum of locus-specific and substrate parameters measured from GPTool with substrate correction (red). The latter fits best the experimental curves. **b**) Radial ensemble MSD curves for *Sox2*- *SCR* (yellow) and *Inter-Down* (black) locus pairs, with the lines showing the median value and the shading the median absolute deviation. Dotted lines show the curves derived from the fitted diffusive parameters. **c**) Bar plot showing the distributions of *α_app_* measurements for the four loci used in this study, compared to the three loci within the HoxA region measured in ref 25. Bars show median values, with the error bars indicating median absolute deviation. **d**) 4 kb resolution Hi-C contact maps derived from the molecular dynamics simulation (left) and the experimental data from ref 60 (right), showing that the simulation recapitulates the major architectural features of the *Sox2* locus. **e**) Violin plots showing the distributions of apparent anomalous exponents measured from the individual movies for *Sox2*, SCR, Inter and Down regions, comparing treatments with DMSO carrier or triptolide. Comparisons are made by Wilcoxon rank sum tests, with *p*-values given.

**Figure S6.**
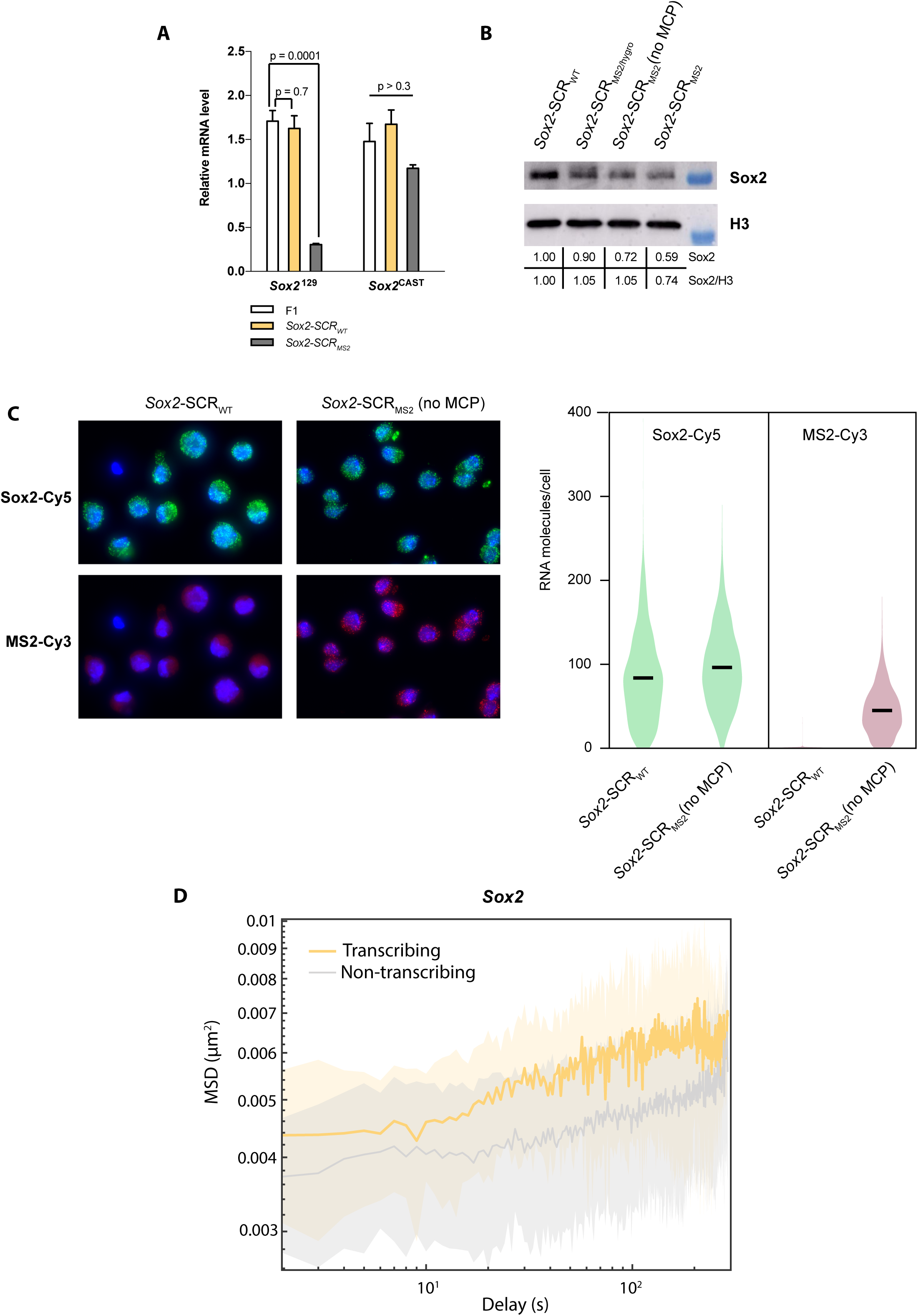
The MS2 system mildly perturbs *Sox2* expression. **a**) Allele-specific qRT-PCR quantification of *Sox2* expression, normalized to *SDHA*, for F1, *Sox2-SCR_WT_*and *Sox2-SCR_MS2_* cells (*n* >= 3). Error bars show standard deviations of the mean. Comparisons are made by two-tailed t-tests, with *p*- values given. **b**) Western blot for SOX2 protein and histone H3 from *Sox2-SCR_WT_*, *Sox2-SCR_MS2_* and intermediate cell lines (before excision of selectable marker - *Sox2-SCR_MS2/hygro_*; before stable insertion of ParB (OR) constructs and MCP - *Sox2-SCR_MS2_ (no MCP)*). Quantification of SOX2 and SOX2/H3 ratios are given, expressed as proportion of *Sox2-SCR_WT_*. **c**) Representative images (left) and violin plots (right) for single-molecule RNA FISH, assessing expression of *Sox2* (green) and MS2 repeats (red), in *Sox2-SCR_WT_*and *Sox2-SCR_MS2_ (no MCP)* cells. Violin plots show distributions of mRNA counts per cell. Total *Sox2* mRNA levels are relatively similar between the two cell lines, and in *SCR_MS2_ (no MCP)* cells, MS2 mRNA levels are around half that of *Sox2*, consistent with its monoallelic expression. **d**) MSD ensemble curves, corrected for substrate movement and plotted on a double-log scale, comparing *Sox2* dynamics between transcriptionally active and inactive alleles. Lines indicate median values and shading the median absolute deviation.

**Figure S7.**
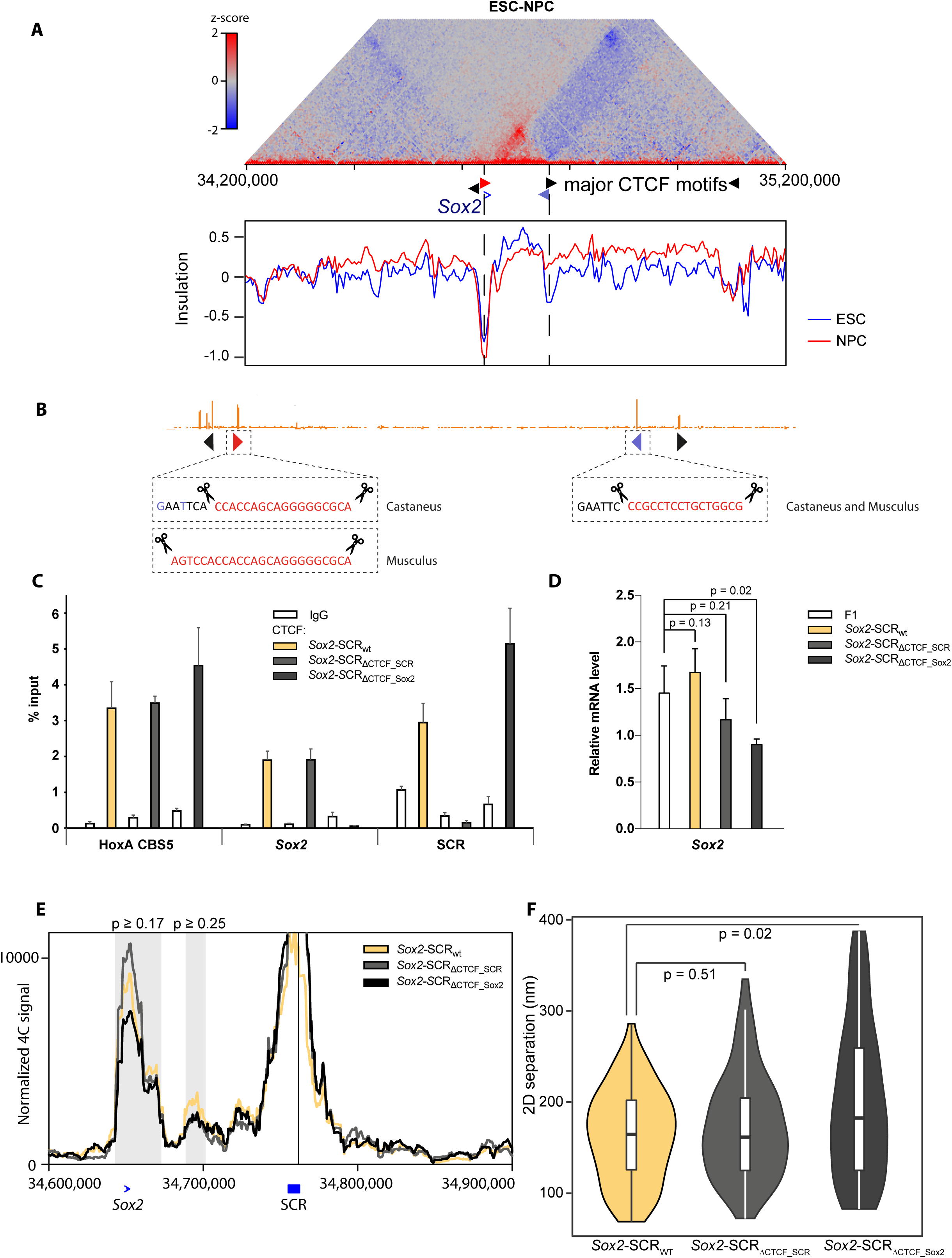
CTCF site deletions at the *Sox2* locus. **a**) Differential Hi-C map, comparing chromatin architecture in ESCs and NPCs around the *Sox2* locus (data taken from ref 60). The red domain indicates a TAD delimited by *Sox2* and SCR which is specific to ESCs. Below are graphs of insulation score computed for the same genomic region, with dotted lines showing the positions of ESC local minima of insulation (i.e. TAD borders) at *Sox2* and SCR. The insulation local minimum at *Sox2* is maintained in NPCs but lost at the SCR. **b**) ESC CTCF ChIP-seq profile around a zoomed in region of the *Sox2* locus, with the orientations of CTCF motifs shown as arrows underneath. The genotypes of the *Sox2-SCR_ΔCTCF_Sox2_* and *Sox2-SCR_ΔCTCF_SCR_*cells are shown underneath. For *Sox2-SCR_ΔCTCF_SCR_*, both alleles have the exact same deletion of 16 nt (orange sequence) at the SCR CTCF motif (blue arrow) and insertion of an *Eco*RI site (black sequence). For *Sox2-SCR_ΔCTCF_Sox2_*, the castaneus allele has deletion of 18 nt at the *Sox2* CTCF motif (red arrow), and mutation of two nucleotides (blue) in the upstream region to generate the *Eco*RI site; the musculus allele has a larger 24 nt deletion at the same motif. **c**) ChIP-qPCR quantification, expressed as percentage of input, of amount of CTCF binding at the HoxA locus CBS5, and *Sox2* and SCR CTCF sites, in *Sox2-SCR_WT_*, *Sox2-SCR_ΔCTCF_Sox2_*and *Sox2-SCR_ΔCTCF_SCR_* cells, compared to mock immunoprecipitation with IgG. CTCF binding is clearly lost at the sites with the motif deletions. **d**) qRT-PCR quantification of *Sox2* expression, normalized to *SDHA*, in F1, *Sox2-SCR_WT_*, *Sox2-SCR_ΔCTCF_Sox2_* and *Sox2-SCR_ΔCTCF_SCR_*cells (*n* >=3). Error bars show standard deviations of the mean. Comparisons of expression with the F1 founder line are made by two-tailed t-tests, with *p*-values given. **e**) Musculus-specific 4C-seq profiles (mean of two replicates) using the SCR as bait are shown *Sox2-SCR_WT_*, *Sox2-SCR_ΔCTCF_Sox2_* and *Sox2-SCR_ΔCTCF_SCR_* cells. The two regions consistently called as interactions are denoted in gray, and the minimum *p*-values for compared interaction scores with *Sox2-SCR_WT_* (two-tailed t-test) are denoted. **f**) Violin plot showing the distributions of median inter-probe distances for *Sox2-SCR_WT_*, *Sox2-SCR_ΔCTCF_Sox2_* and *Sox2-SCR_ΔCTCF_SCR_* cells. Comparisons are made by Wilcoxon rank sum tests, with *p*-values given.

**Figure S8.**
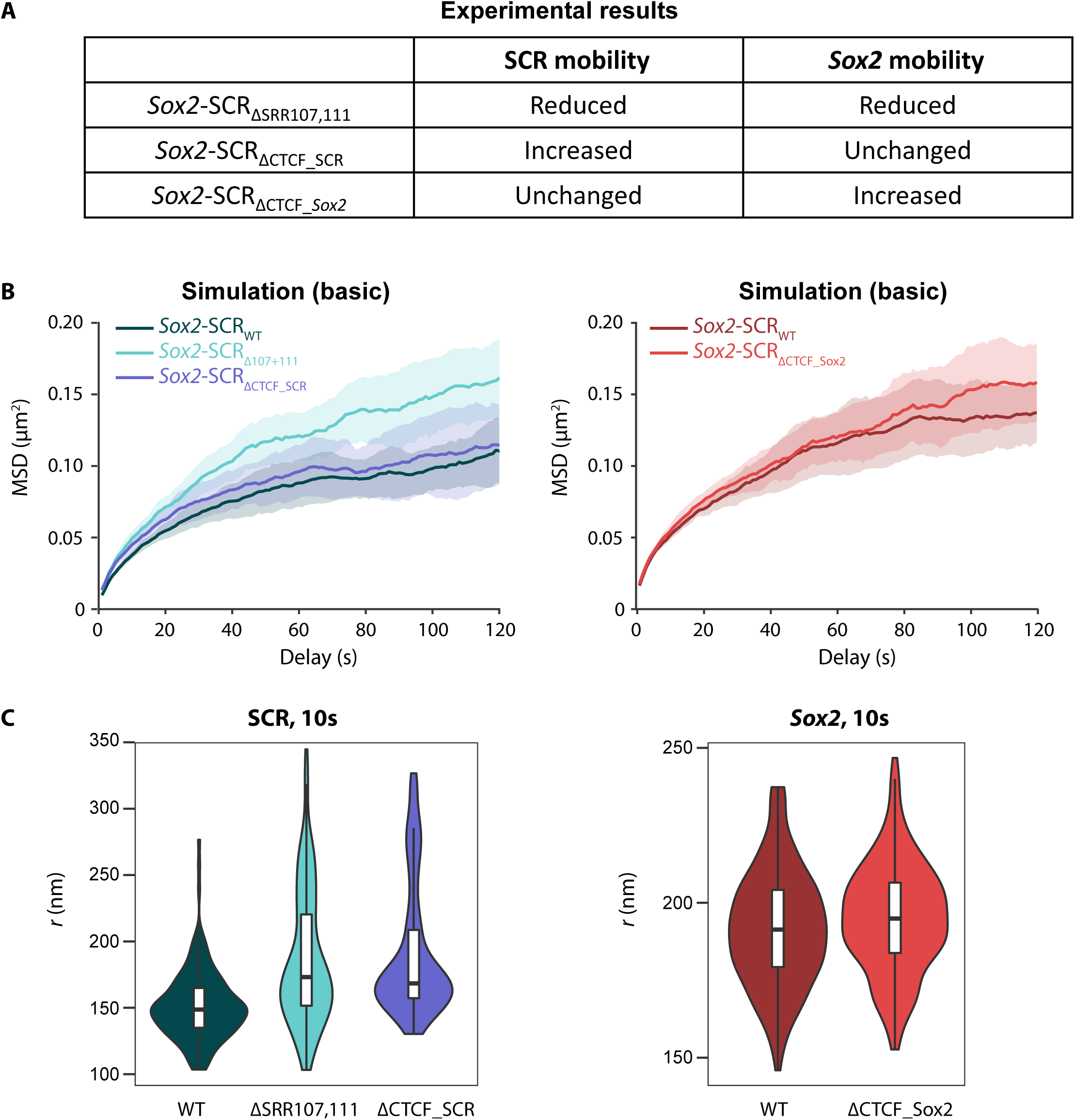
The original physical model poorly predicts effects of genetic perturbations on chromatin mobilities at the *Sox2* locus. **a**) Table summarizing the measured effects on SCR and *Sox2* mobility on the different genetic perturbations covered in this study, compared to wild-type. **b**) Ensemble MSD curves for the SCR (left) and *Sox2* (right), derived from molecular dynamics simulations of the Sox2 locus in wild-type or SCR mutant conditions. Lines show median values and shading indicates the median absolute deviation. **c**) Violin plots showing distributions of apparent explored radii of the SCR (left) and *Sox2* (right) after 10 s, derived from the same simulations as **b**. These simulations predict increased SCR mobility on deletion of SRR107 and 111, not the experimentally measured decrease.

**Figure S9.**
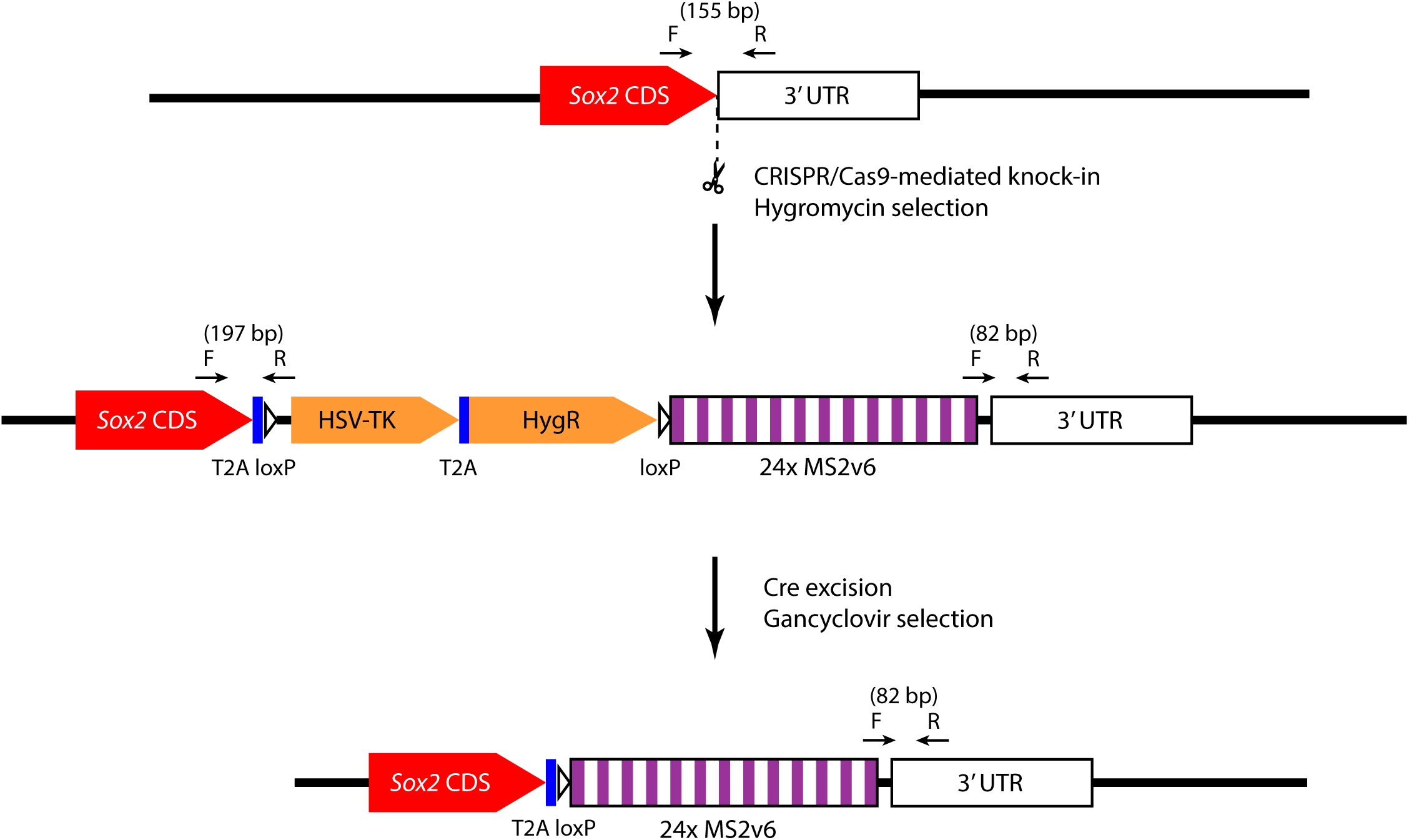
Schematic for insertion of MS2 repeats in generation of *Sox2-SCR_MS2_* line. Construct containing 24x MS2v6 repeats and the positive-negative selection marker (HSV-TK/HygroR) is knocked in directly adjacent to the STOP codon of *Sox2* with CRISPR/Cas9-mediated homologous recombination, and clones with insertions are selected with hygromycin. Inclusion of binding sequences for primers (F and R) flanking the *Sox2* STOP codon in the knocked-in construct allows one PCR reaction to distinguish wild-type (155 bp product) from inserted (82 bp and 197 bp products) alleles. Heterozygous clones, producing all three PCR products are then screened further with Sanger sequencing to determine in which allele the MS2 repeats were inserted. The selectable marker, which may affect *Sox2* transcription, is then excised by transfection with a Cre-expressing plasmid, selecting clones with gancyclovir treatment, and screening for expected loss of 197 bp PCR product. Strategy adapted from ref 71.

**Table S1. Chromatin dynamics measured by double-label ANCHOR experiments.** Details for each double-label ANCHOR trajectory used in this study, giving the experimental condition, individual movie name, median inter-probe distance, the identities of the two channels, and their measured *D_α_app_* and *α_app_* values.

Provided as separate Excel sheet.

**Table S2. Chromatin dynamics measured by triple-label ANCHOR experiments.** Details for each ANCHOR trajectory used in this study from the *Sox2-SCR_MS2_* line, giving the experimental condition (transcribed or non-transcribed), individual movie name, numbers of frames used in inferring the apparent diffusive parameters, median inter-probe distance, the identities of the two ANCHOR channels, and their measured *D_α_app_* and *α_app_* values.

Provided as separate Excel sheet.

**Table S3.**
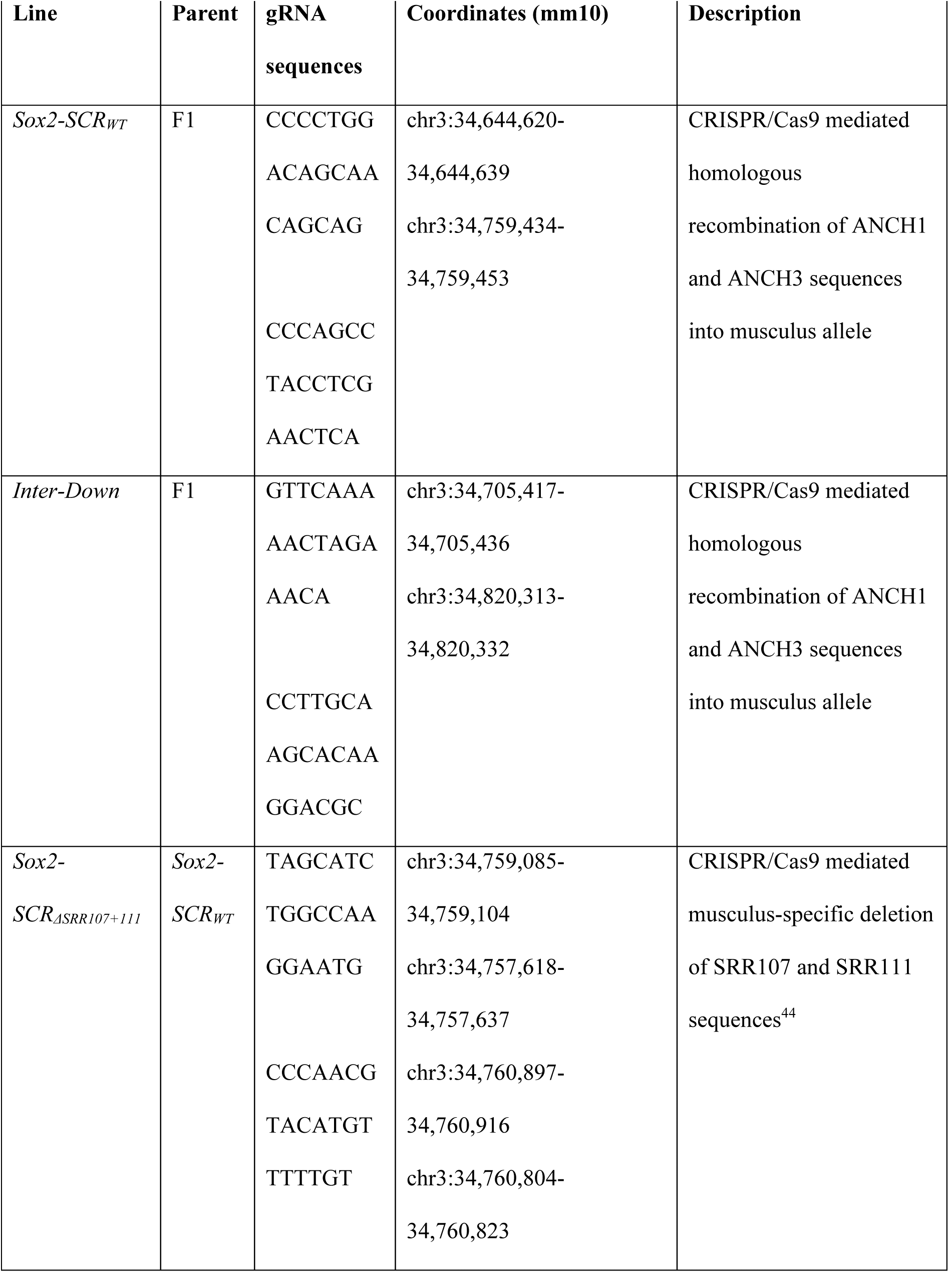
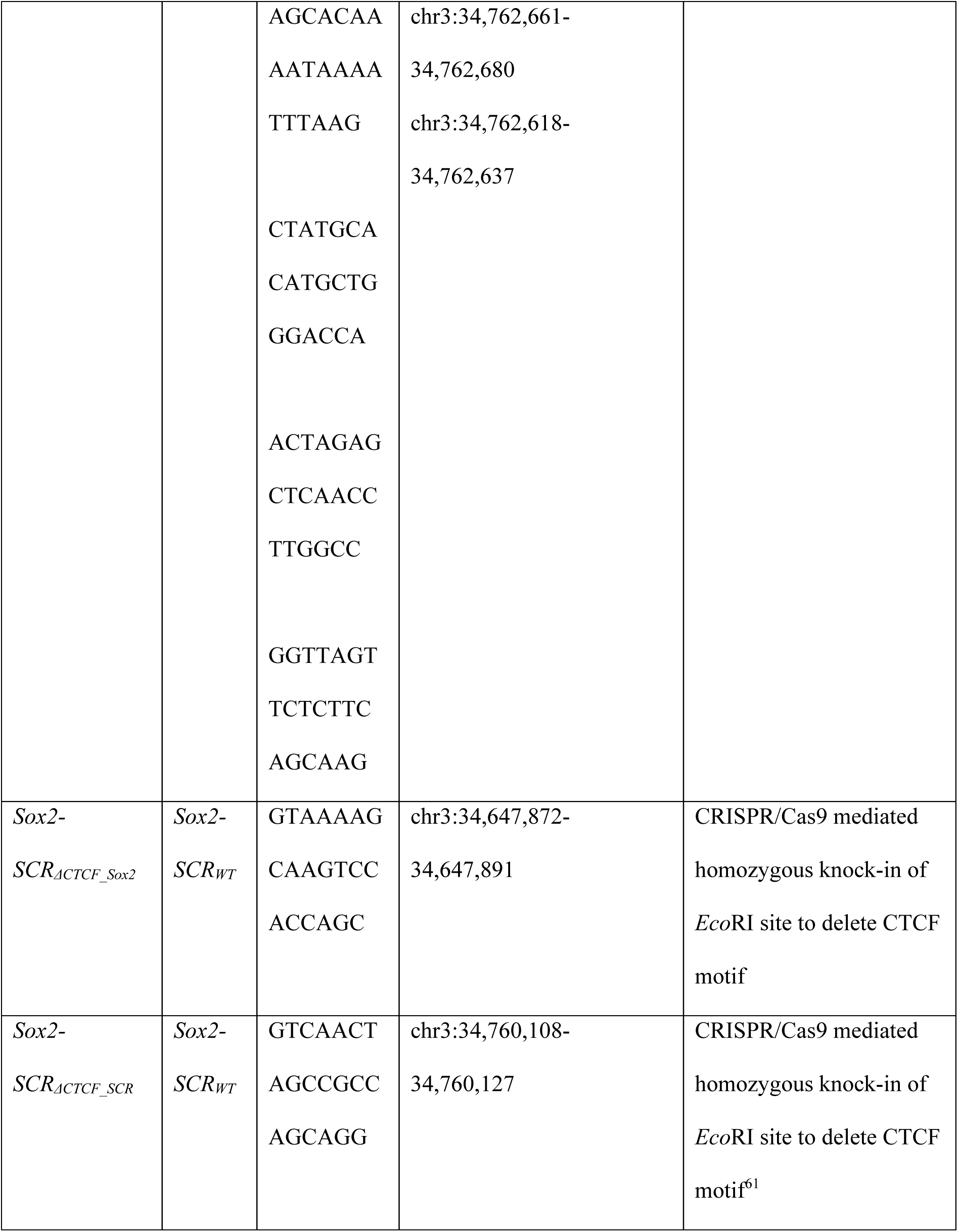
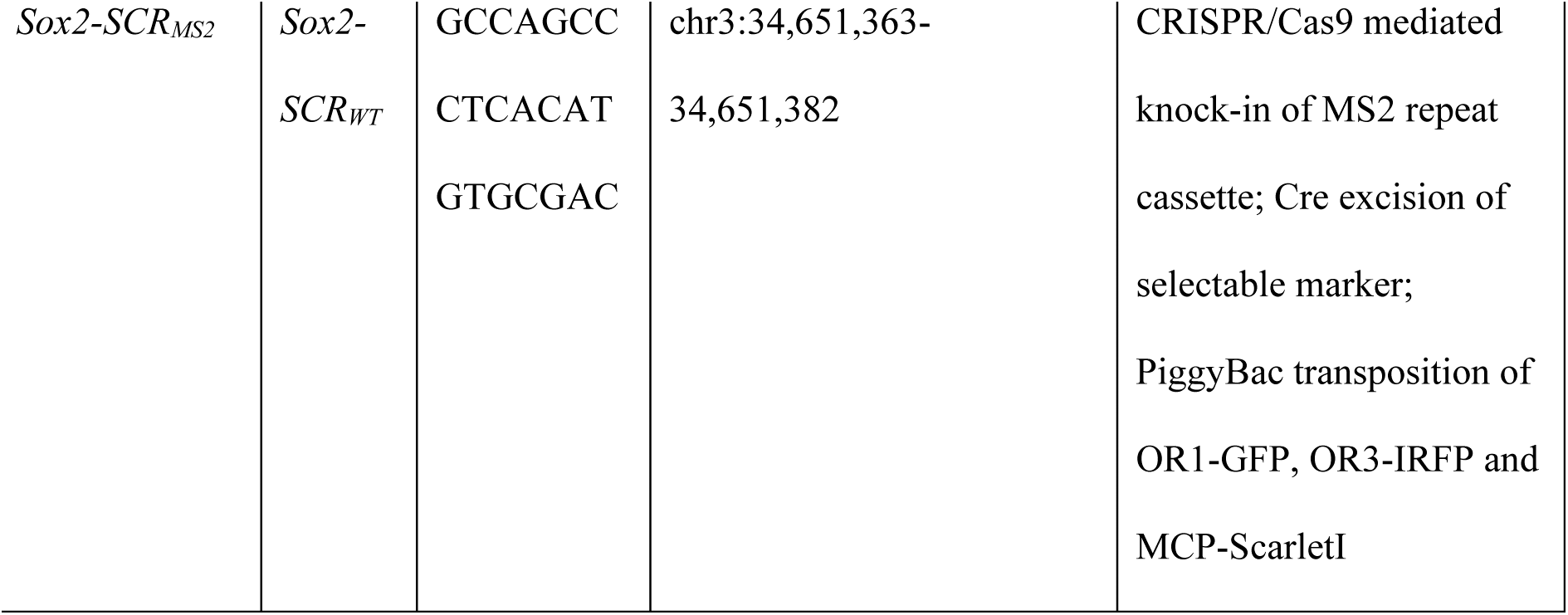
Cell lines generated in this study. Descriptions of the parental lines and gRNA sequences used to make the cell lines used in this study.

**Table S4.**
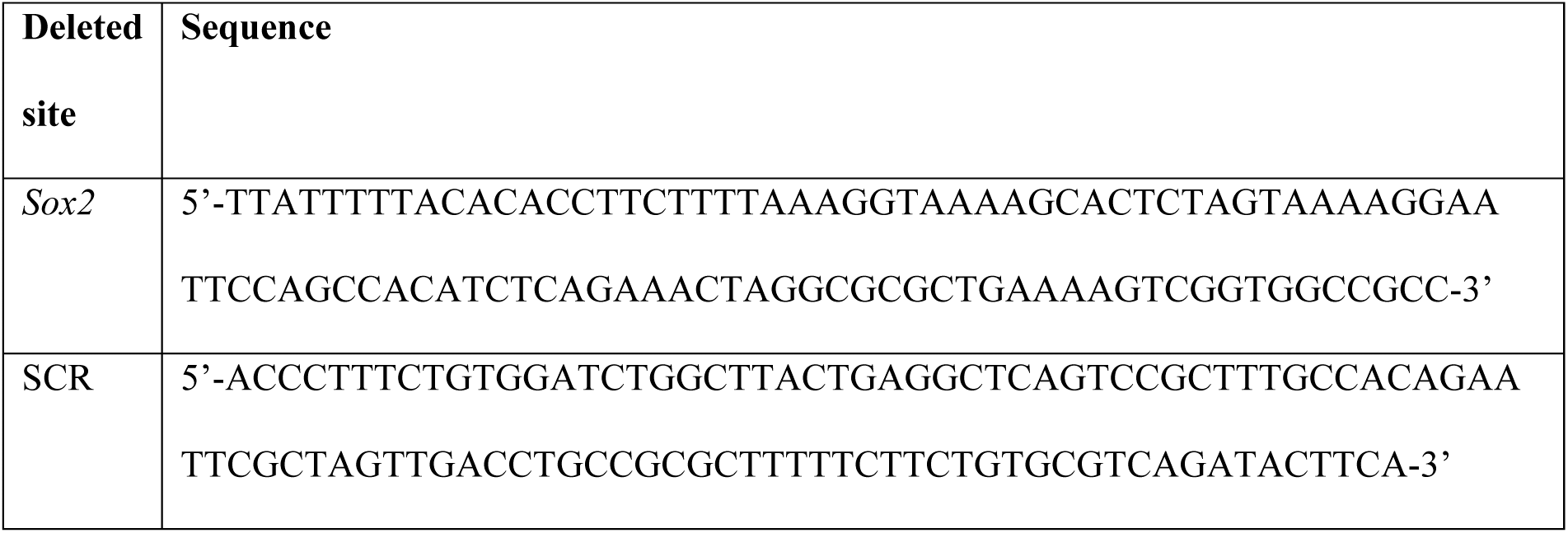
Sequences of repair cassettes used to delete CTCF motifs.

**Table S5.**
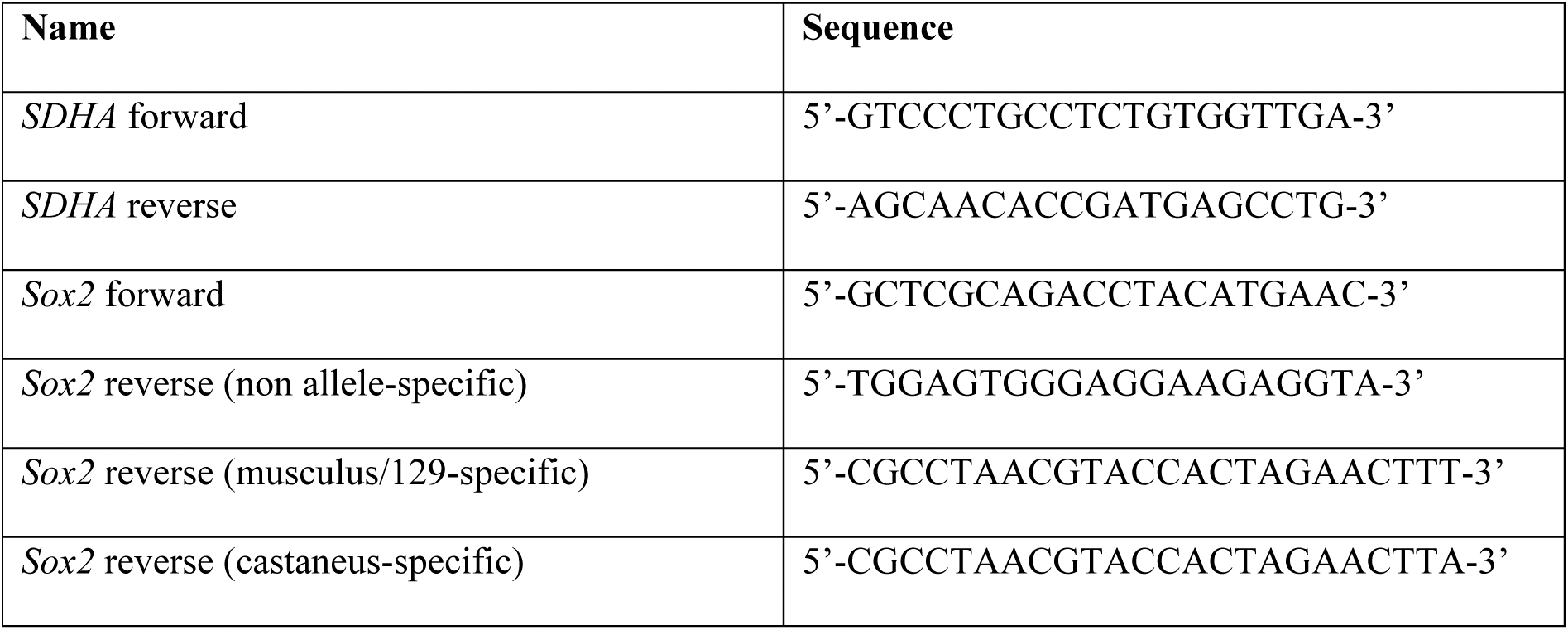
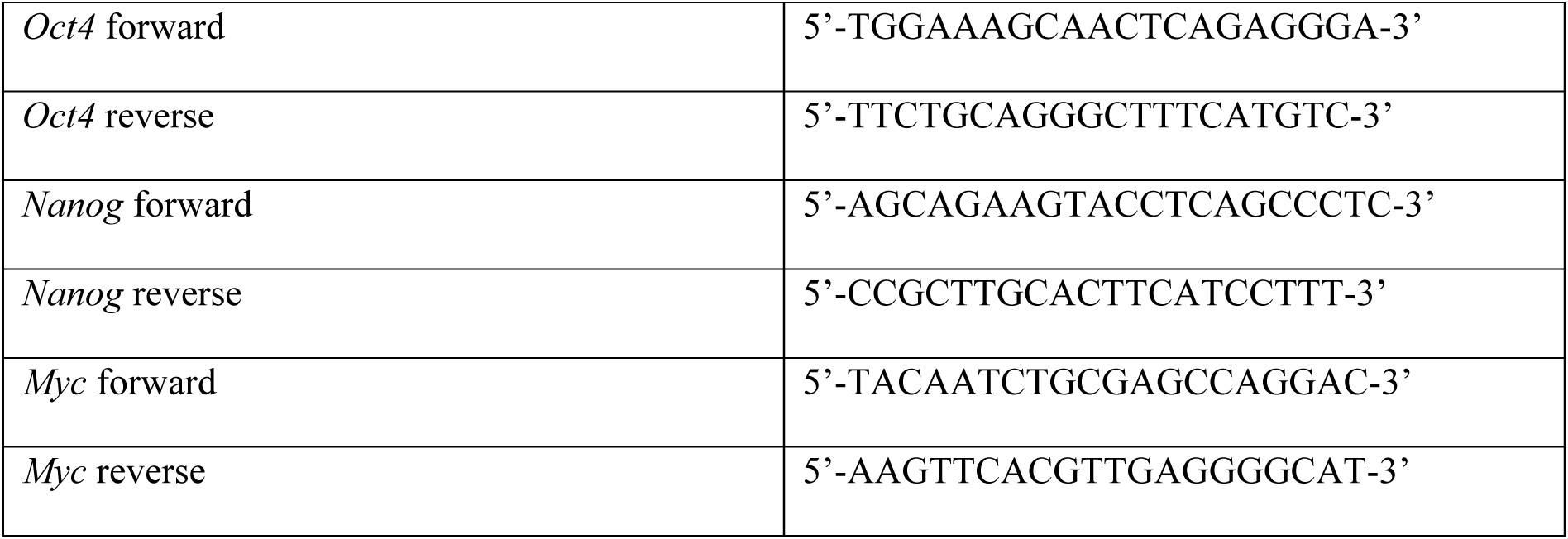
qRT-PCR primer sequences.

**Table S6.**
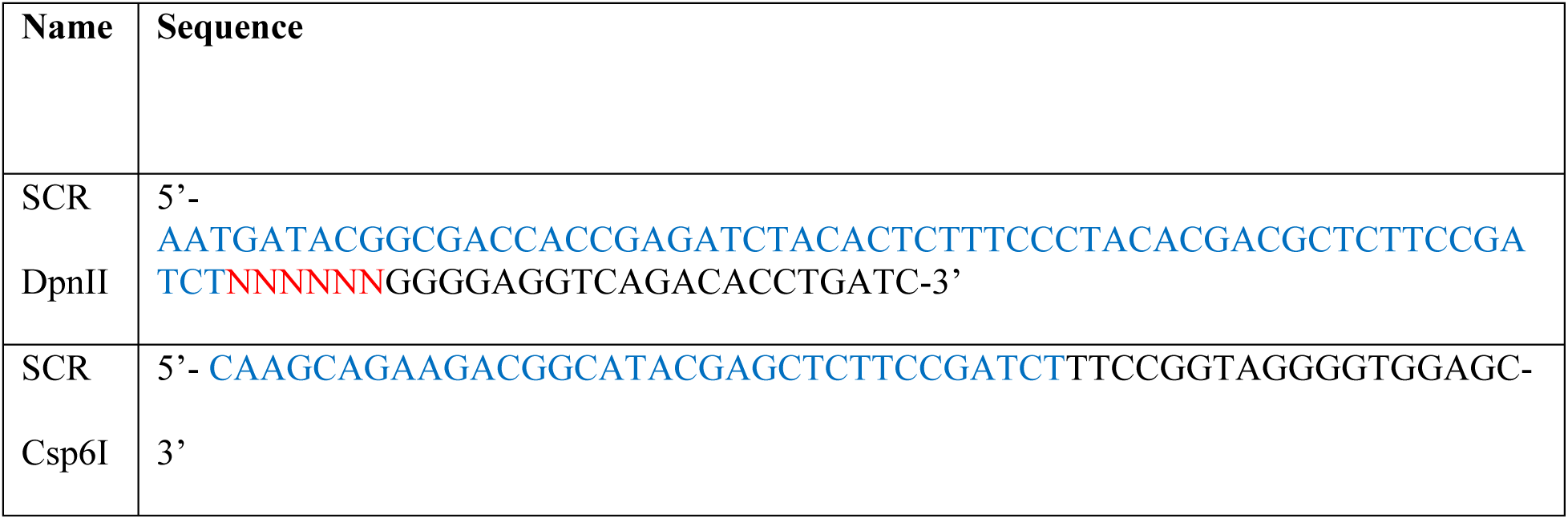
4C-seq primer sequences. Blue denotes Illumina adapter sequence for high-throughput sequencing. Red denotes position of 6-nucleotide barcodes, used to multiplex 4C samples for sequencing.

**Table S7.**
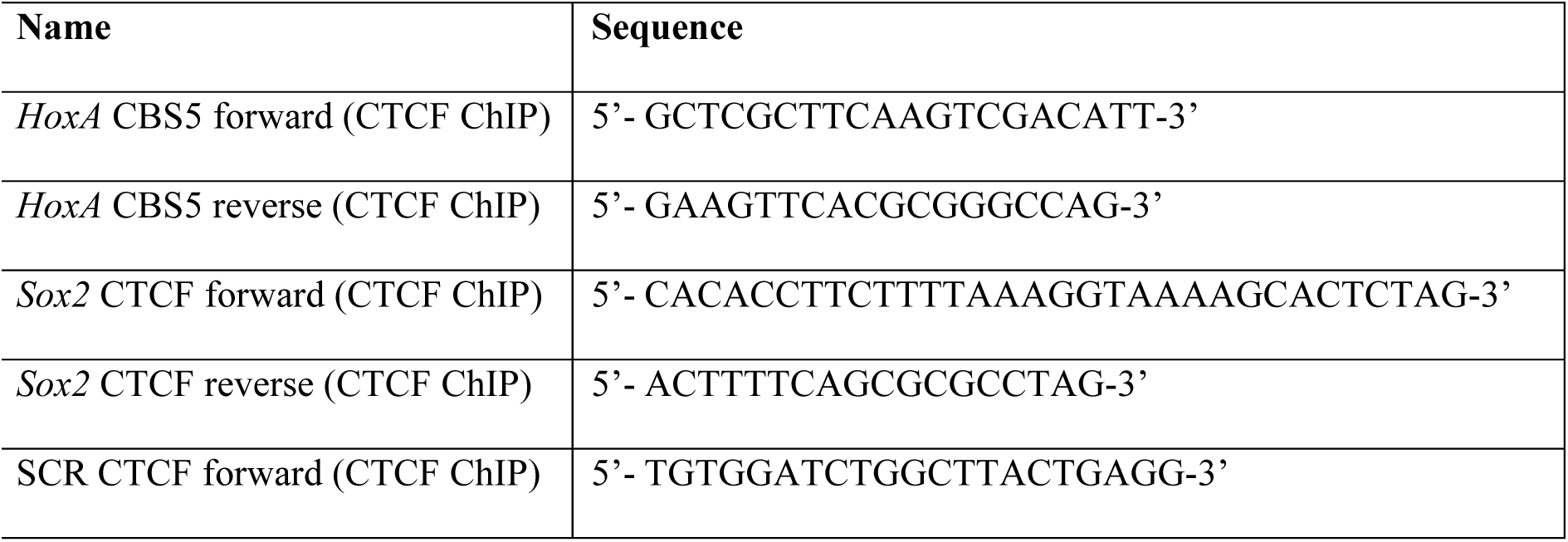
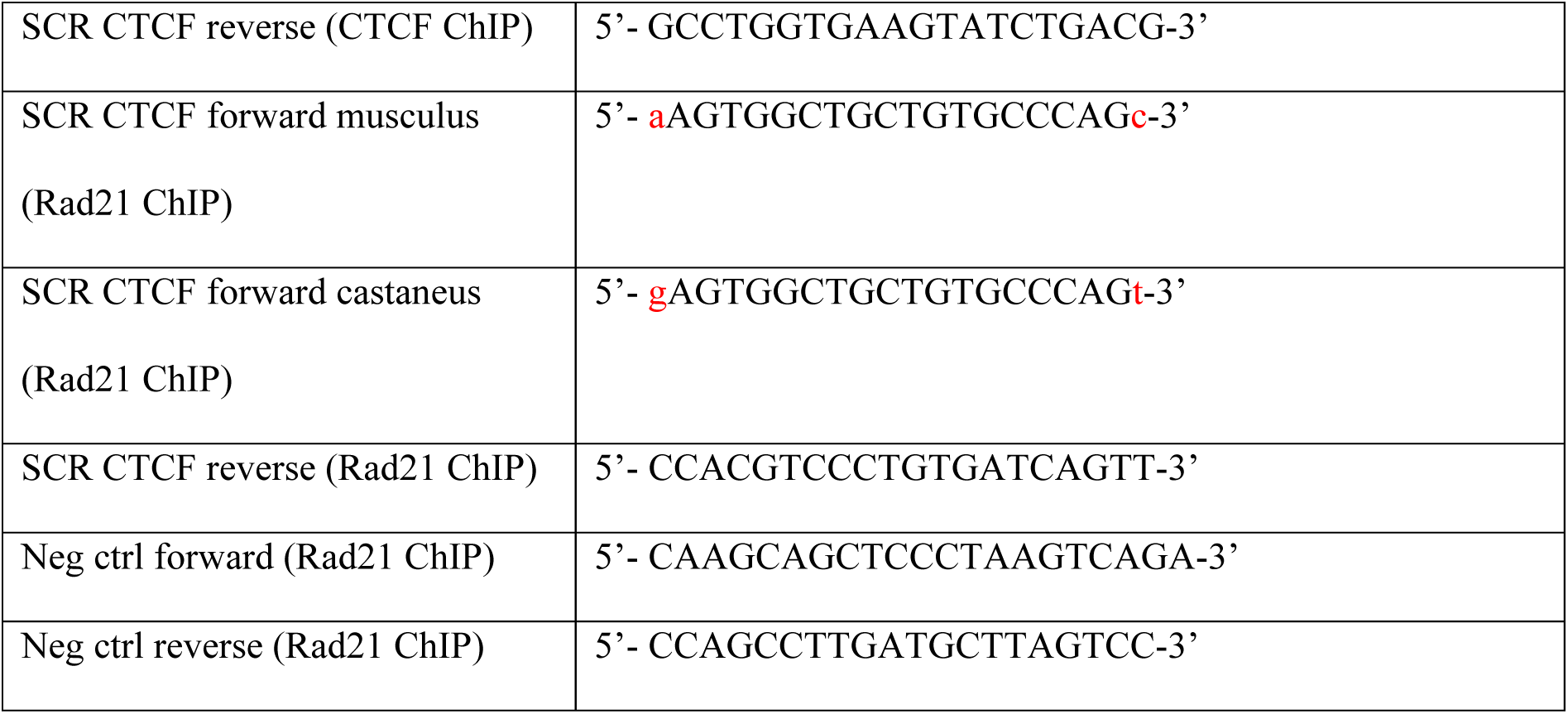
ChIP-qPCR primer sequences. Red denotes allele-specific nucleotides.

## Notes

### Competing Interest Statement

The authors have declared no competing interest.

